# The nascent RNA binding complex SFiNX licenses piRNA-guided heterochromatin formation

**DOI:** 10.1101/609693

**Authors:** Julia Batki, Jakob Schnabl, Juncheng Wang, Dominik Handler, Veselin I. Andreev, Christian E. Stieger, Maria Novatchkova, Lisa Lampersberger, Kotryna Kauneckaite, Wei Xie, Karl Mechtler, Dinshaw J. Patel, Julius Brennecke

**Affiliations:** Institute of Molecular Biotechnology of the Austrian Academy of Sciences (IMBA) Vienna BioCenter (VBC), Dr. Bohrgasse 3, 1030 Vienna, Austria; Structural Biology Program, Memorial Sloan Kettering Cancer Center, New York, NY 10065, USA; Institute of Molecular Pathology (IMP) Vienna BioCenter (VBC), 1030 Vienna, Austria

## Abstract

The PIWI-interacting RNA (piRNA) pathway protects animal genome integrity in part through establishing repressive heterochromatin at transposon loci. Silencing requires piRNA-guided targeting of nuclear PIWI proteins to nascent transposon transcripts, yet the subsequent molecular events are not understood. Here, we identify SFiNX (Silencing Factor interacting Nuclear eXport variant), an interdependent protein complex required for Piwi-mediated co-transcriptional silencing in *Drosophila*. SFiNX consists of Nxf2-Nxt1, a gonad-specific variant of the heterodimeric mRNA export receptor Nxf1-Nxt1, and the Piwi-associated protein Panoramix. SFiNX mutant flies are sterile and exhibit transposon de-repression because piRNA-loaded Piwi is unable to establish heterochromatin. Within SFiNX, Panoramix recruits the heterochromatin effectors, while the RNA binding Nxf2 protein licenses co-transcriptional silencing. Our data reveal how Nxf2 evolved from an RNA transport receptor into a co-transcriptional silencing factor. Thus, NXF-variants, which are abundant in metazoans, can have diverse molecular functions and might have been co-opted for host genome defense more broadly.

## INTRODUCTION

Eukaryotic cells establish heterochromatin at genomic repeats and transposon insertions to suppress transcription and ectopic recombination (Fedoroff, 2012; Slotkin and Martienssen, 2007). One strategy to confer sequence specificity to this process is via repressor proteins (e.g. KRAB-type zinc finger repressors in tetrapods) that bind defined DNA motifs and recruit heterochromatin inducing factors (Yang et al., 2017). A second strategy for sequence-specific heterochromatin formation builds on nuclear small RNAs (Castel and Martienssen, 2013; Grewal, 2010; Holoch and Moazed, 2015). These ~20-30nt long regulatory RNAs guide Argonaute proteins to complementary sequences in nascent target transcripts, which are still attached to chromatin via transcribing RNA polymerases (Shimada et al., 2016). Binding of nuclear Argonautes to nascent transcripts leads to transcriptional repression and heterochromatin formation. As Argonaute recruitment requires the nascent target RNA, this process is defined as ‘co-transcriptional silencing’. Besides impacting chromatin and transcription, nuclear small RNA pathways have also been linked to the co-transcriptional processes of splicing, RNA quality control and turnover (Dumesic et al., 2013; Reyes-Turcu et al., 2011; Teixeira et al., 2017). Together, this hints at complex molecular connections between nuclear Argonautes, the nascent target RNA, and chromatin. Most of our knowledge on nuclear small RNA-guided silencing is based on work in fission yeast and plants. How Argonautes orchestrate c-transcriptional silencing in animals is, however, not understood.

The principal nuclear Argonaute pathway in animals is the PIWI-interacting small RNA (piRNA) pathway (Czech et al., 2018; Ozata et al., 2018). It acts preferentially in gonads and safeguards the integrity of the germline genome. In *Drosophila melanogaster*, a single nuclear Argonaute protein (Piwi) orchestrates co-transcriptional silencing and heterochromatin formation at hundreds of transposon insertions throughout the genome (Le Thomas et al., 2013; Rozhkov et al., 2013; Sienski et al., 2012; Wang and Elgin, 2011). Although transposons contain strong promoters and enhancers, binding of Piwi to nascent transposon transcripts effectively suppresses their transcription, resulting in up to several hundred-fold reductions in steady state RNA levels. How Piwi orchestrates co-transcriptional silencing at the molecular level is poorly understood. Genetic studies identified three piRNA pathway-specific proteins, Maelstrom, Asterix/Gtsf1, and Panoramix/Silencio, as essential co-factors for Piwi-mediated silencing (Donertas et al., 2013; Muerdter et al., 2013; Ohtani et al., 2013; Sienski et al., 2015; Sienski et al., 2012; Yu et al., 2015). In the absence of any of these proteins, Piwi is abundantly expressed, localizes to the nucleus, is loaded with transposon-targeting piRNAs, but is incapable of target silencing. Among these three Piwi co-factors, only Panoramix is capable of inducing co-transcriptional silencing and heterochromatin formation if targeted to a nascent RNA through aptamer-based tethering. Silencing via tethered Panoramix is independent of Piwi but requires the H3K9 methyltransferase Eggless/SetDB1, the H3K4 demethylase Su(var)3-3/Lsd1, and the H3K9me2/3 reader protein HP1 (Sienski et al., 2015; Yu et al., 2015). This places Panoramix downstream of Piwi and upstream of the cellular heterochromatin machinery. How Panoramix, which does not resemble any known protein, is connected to the nascent RNA, or to chromatin effectors and chromatin itself, is unknown.

Here, we show that Panoramix functions in the context of an interdependent protein complex with Nxf2-Nxt1/p15, a variant of the highly conserved Nxf1/Tap-Nxt1/p15 heterodimer. From budding yeast to humans, Nxf1/Tap-Nxt1/p15 is the principal nuclear mRNA export receptor that mediates the translocation of export-competent mRNAs through nuclear pore complexes into the cytoplasm (Cullen, 2000; Kohler and Hurt, 2007). Nxf2 is one of the three <u>n</u>uclear RNA e<u>x</u>port factor (NXF) variants in *Drosophila melanogaster*, but no mRNA export function could be attributed to it (Herold et al., 2001; Herold et al., 2003). Using genetics, biochemistry and structural biology, we reveal an unexpected, RNA-export independent role for Nxf2 in piRNA guided co-transcriptional silencing of transposons. Our findings provide a functional link between the genome-transposon conflict and NXF-variants, which have largely unknown function in animals.

## RESULTS

### The Nxf2-Nxt1/p15 heterodimer interacts with Panoramix

To elucidate the molecular function of Panoramix, we determined its protein interactors in cultured ovarian somatic cells (OSCs), which express a nuclear Piwi/piRNA pathway (Niki et al., 2006; Saito et al., 2009). We isolated Panoramix via immuno-precipitation (IP) from nuclear lysate of a clonal OSC line expressing FLAG-tagged Panoramix and identified co-eluting proteins by quantitative mass spectrometry. The most prominent interactors were nuclear RNA export factor 2 (Nxf2), the mRNA export co-factor Nxt1/p15, and eIF-4B (Fig. 1a left; Table S1). Among those, Nxf2 and Nxt1/p15 were also identified in genetic transposon de-repression screens (Czech et al., 2013; Handler et al., 2013; Muerdter et al., 2013). We confirmed the interaction between Panoramix, Nxf2 and Nxt1/p15 using reciprocal co-IP mass-spectrometry with FLAG-tagged Nxf2 as bait (Fig. 1a right; Table S1). In both experiments, peptide levels for bait and interactors were in a similar range, suggesting that Panoramix, Nxf2, and Nxt1/p15 form a stable protein complex (see below; Supplementary Fig. 1a). In comparison, the previously identified Panoramix-interactor Piwi (Sienski et al., 2015; Yu et al., 2015) was only ~2-fold enriched and Piwi peptide levels in the IP eluates were ~20-fold lower than the other interactors (Supplementary Fig. 1a, Table S1), indicating a transient or sub-stoichiometric association between Piwi and Panoramix or Nxf2. A co-IP experiment using a monoclonal antibody against Panoramix confirmed these findings for the endogenous Panoramix, Nxf2 and Piwi proteins (Fig. 1b).

**Figure 1.**
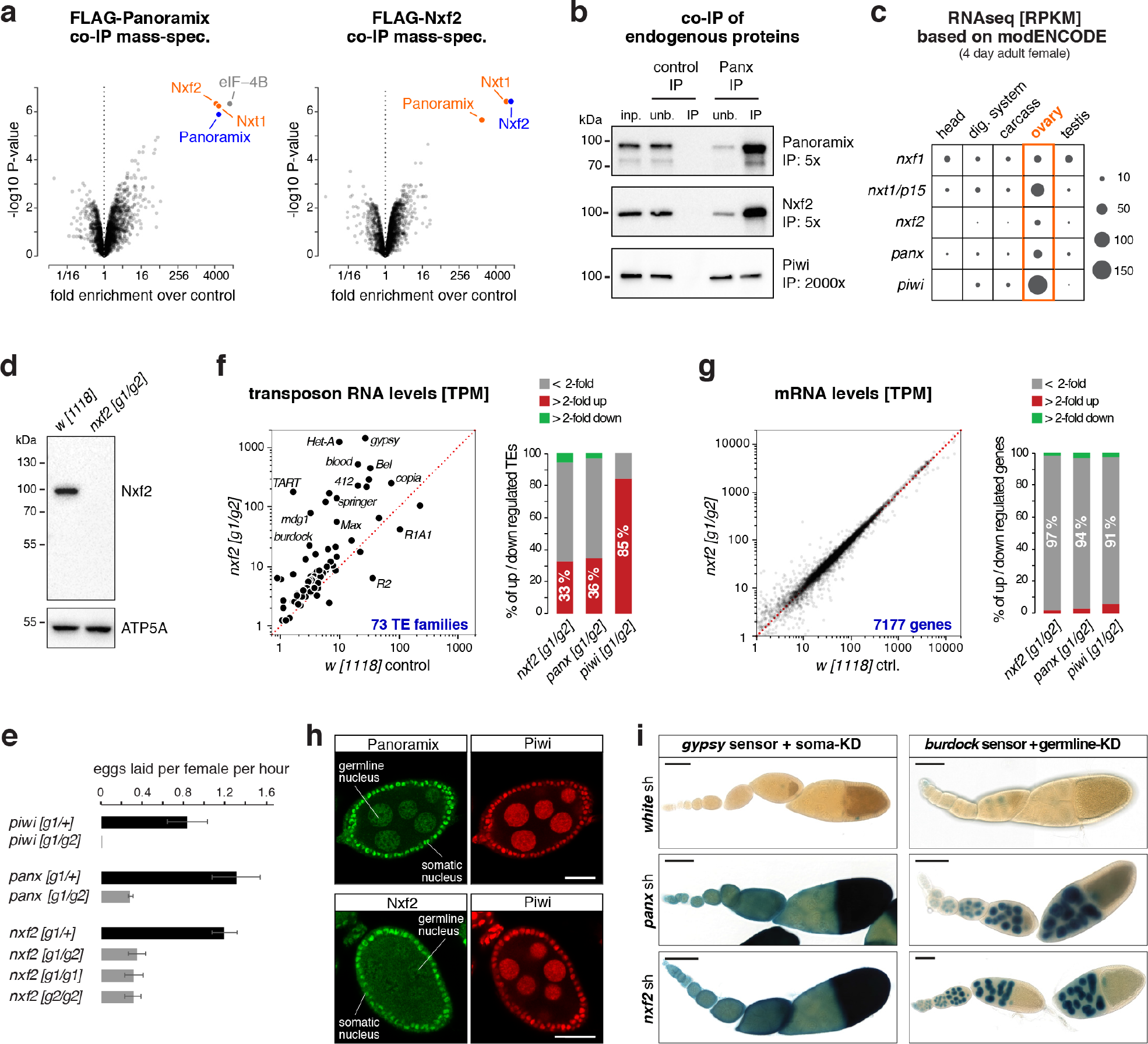
The Nxf2-Nxt1/p15 heterodimer interacts with Panoramix. **a**, Volcano plots showing fold enrichments of proteins determined by quantitative mass spectrometry in FLAG-Panoramix (left) or FLAG-Nxf2 (right) co-immuno-precipitates versus control (n = 3). Experiments were performed with OSC cell lines stably expressing cDNA constructs encoding tagged Panoramix or Nxf2 (wildtype OSCs served as control). **b**, Western blot analysis of co-immunoprecipitation of endogenous Panoramix from nuclear OSC lysate. Relative amount loaded in immunoprecipitation (IP) compared to input (inp.) is indicated for each protein. Mouse IgGs were used as negative control. **c**, Shown are expression levels of indicated genes based on modENCODE mRNA-seq data in different adult tissues. **d**, Western blot analysis of ovarian lysates (genotypes indicated) with monoclonal anti-Nxf2 antibody. ATP5A served as loading control. **e**, Bar plots showing average egg-laying rates of females with indicated genotype (n=3; error bars indicate SD). None of the eggs laid by *panoramix* or *nxf2* mutant females developed into larvae, indicating complete female sterility. **f, g** Left: Scatter plots showing steady state ovarian RNA levels (transcripts per million) of genes (**f**) and transposons (**g**) in *nxf2* mutants compared to control. Right: Bar diagrams indicate differentially expressed genes or transposons in indicated genotypes. **h**, Confocal images of egg chambers (scale bar: 20 μm) from wild-type flies co-stained for Panoramix (top) or Nxf2 (bottom) with Piwi. For specificity of antibodies see Sup. Fig. 1f. **i**, Shown are X-Gal stained ovarioles (scale bar: 0.1mm) from flies expressing *gypsy*-lacZ (soma sensor; left) or *burdock*-lacZ (germline sensor; right) plus indicated knockdown (KD) constructs (sh-lines).

Nxf2 belongs to the NXF protein family, which in *Drosophila* is composed of the principal mRNA export receptor Nxf1, and the NXF variants Nxf2, Nxf3, and Nxf4 (Herold et al., 2001; Herold et al., 2000). Like *piwi* and *panoramix*, *nxf2* is expressed predominantly in ovaries (Fig. 1c) (Brown et al., 2014). To follow up on the connection between Panoramix and Nxf2, we generated *nxf2* null mutant flies (Fig. 1d; Supplementary Fig. 1b). In contrast to *nxf1* and *nxt1/p15* mutants, which are lethal (Caporilli et al., 2013; Wilkie et al., 2001), *nxf2* mutants were viable and developed gonads (Supplementary Fig. 1c). However, as *panoramix* mutants, *nxf2* mutant females were sterile; compared to control flies, they laid fewer eggs (Fig. 1e) and none of these developed into a larva. To investigate whether the sterility of *nxf2* mutants is linked to defects in transposon silencing, we sequenced total ovarian RNA from *nxf2* mutants and control flies. Similar to *panoramix* or *piwi* mutants, *nxf2* mutants expressed strongly elevated levels of several transposon families (30%; TPM>5) while only very few endogenous mRNAs exhibited changes in their levels (Fig. 1f, g; Supplementary Fig. 1d, e; Table S2). Among the de-silenced transposons were germline-specific (e.g. *HeT-A*, *burdock*) and soma-specific (e.g. *gypsy*, *mdg1*) elements, indicating that Nxf2 is required for transposon silencing in both ovarian tissues. In support of this, Nxf2 was expressed, like Panoramix, in ovarian somatic and germline cells (Fig. 1h; Supplementary Fig. 1f) and RNAi mediated depletion of Nxf2 specifically in the ovarian soma or germline resulted in de-repression of cell-type specific transposon reporters (Fig. 1i). Taken together, Nxf2 interacts with Panoramix, and is required for fertility and transposon silencing during oogenesis.

### Nxf2 is required for Piwi-mediated heterochromatin formation

Given its interaction with Panoramix, we hypothesized that Nxf2 is required for Piwi to induce co-transcriptional silencing. If so, loss of Nxf2 should not affect piRNA biogenesis, but would result in piRNA-loaded Piwi being unable to specify heterochromatin at target loci (Sienski et al., 2015; Yu et al., 2015). Indeed, Piwi levels and localization were unchanged in *nxf2* mutants (Supplementary Fig. 2a), indicative of intact piRNA biogenesis and loading. Furthermore, sequencing of small RNAs from Nxf2-depleted OSCs, revealed that Nxf2, similar to Panoramix, is not required for piRNA production (Fig. 2a, b). We obtained similar results when we depleted Nxf2 specifically in the ovarian germline via transgenic RNAi (Supplementary Fig. 2b, c). Thus, irrespective of tissue and genomic origin, loss of Nxf2 does not impact piRNA biogenesis.

**Figure 2.**
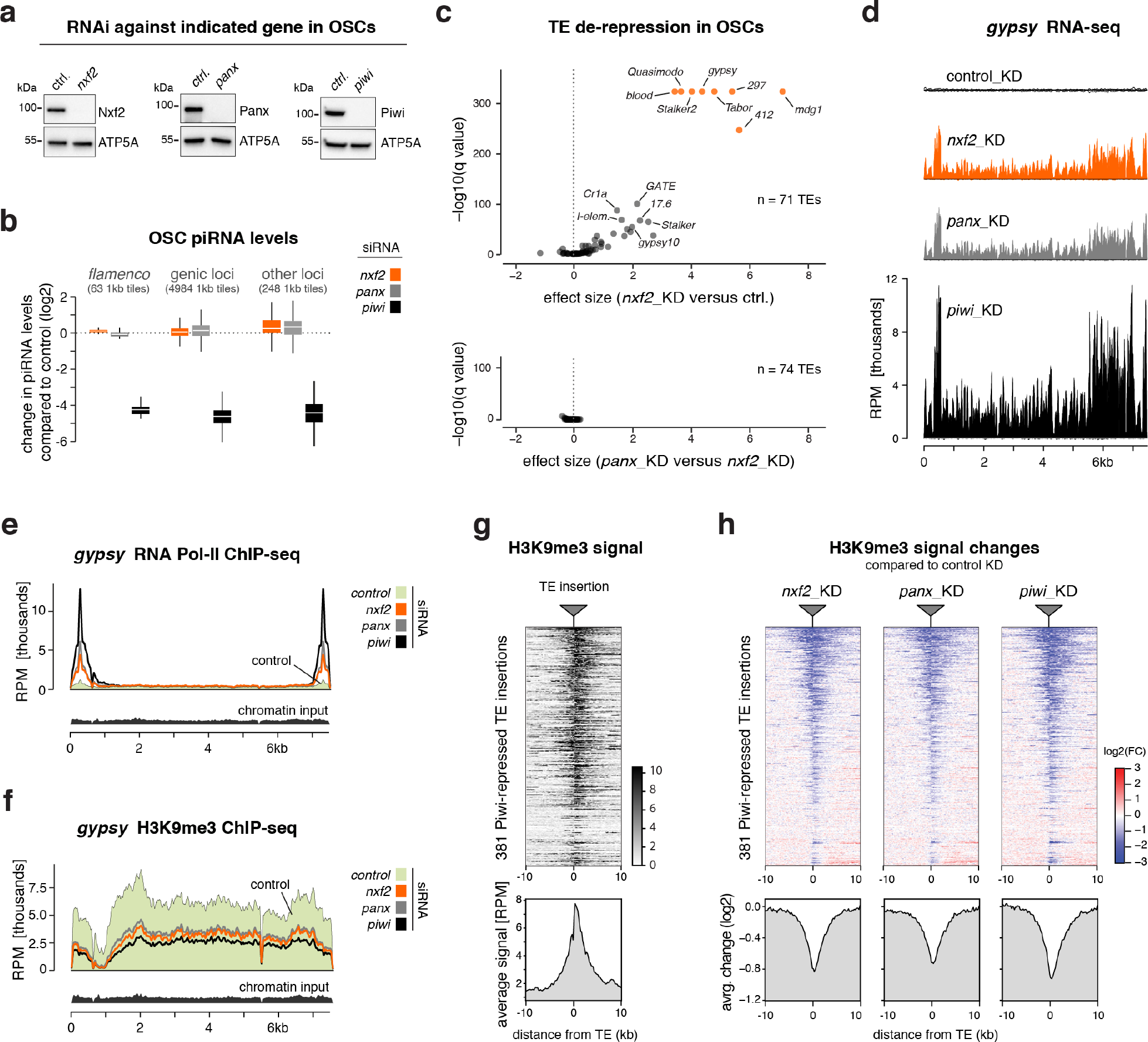
Nxf2 is required for Piwi-mediated heterochromatin formation. **a**, Western blots showing knockdown efficiencies of indicated proteins in OSCs. ATP5A served as loading control. **b**, Box plot showing fold change (compared to control knockdown) of piRNA levels in OSCs depleted of indicated genes (box plots indicate median, first and third quartiles (box), whiskers show 1.5× interquartile range; outliers were omitted). **c**, Volcano plots showing changes (as effect size) in steady state transposon levels from RNA-seq experiments (n=3; upper plot: *nxf2* knockdown versus control, lower plot: *panoramix* versus *nxf2* knockdowns). **d**, Density profiles showing normalized reads from OSC RNA-seq experiments mapping to the *gypsy* transposon consensus sequence (knockdowns indicated). **e, f**, Density profiles showing normalized reads from OSC RNA Pol II ChIP-seq (**e**), or H3K9me3 ChIP-seq (**f**) experiments mapping to the *gypsy* transposon consensus sequence (knockdowns indicated). **g**, Heatmap (top) and metaplot (bottom) showing H3K9me3 levels within the 20kb region flanking all 381 Piwi-repressed euchromatic transposon insertions in the OSC genome (insertions sorted for average H3K9me3 signal per window). **h**, Heatmaps (top) and metaplots (bottom) showing control-normalized log2-fold changes in H3K9me3 levels at genomic regions surrounding transposon insertions defined in Fig. 2g (sorting as in g; gene knockdowns indicated).

To ask whether Nxf2 is required for piRNA-guided co-transcriptional silencing, we turned to OSCs. Here, loss of Piwi results in up to hundred-fold elevated RNA levels of a subset of LTR-retrotransposons due to increased transcription accompanied by loss of the heterochromatic H3K9me3 mark (Supplementary Fig. 2d) (Sienski et al., 2012). In Nxf2-depleted OSCs, all piRNA pathway-repressed transposons were de-silenced despite normal piRNA levels (Fig. 2c top, 2d; Table S3). The extent of de-repression was virtually indistinguishable to that seen in Panoramix-depleted cells (Fig. 2c bottom, 2d). Consistent with a role of Nxf2 in co-transcriptional silencing, ChIP-seq experiments revealed that loss of Nxf2 resulted in increased RNA Polymerase II occupancy and reduced H3K9me3 levels for piRNA-targeted transposons like *gypsy*, or *mdg1* (Fig. 2e, f; Supplementary Fig. 2e). In contrast, transposons not under piRNA control in OSCs (e.g. *burdock, F*-element) showed no such changes (Supplementary Fig. 2f, g).

To assess these effects at the level of individual genomic loci, we examined the euchromatic insertion sites of Piwi-repressed transposons in the OSC genome. At these stand-alone transposon insertions, H3K9me3 marks spread into flanking genomic regions, to which sequencing reads could be mapped unambiguously (Fig. 2g) (Sienski et al., 2012). Focusing on the ~380 Piwi-silenced transposon insertions revealed that Nxf2 loss phenocopies the decrease in H3K9me3 levels seen in cells depleted of Panoramix or Piwi (Fig. 2h). Piwi-independent H3K9me3 domains instead were unaffected in Nxf2-depleted cells (Supplementary Fig. 2h).

Several dozen piRNA-repressed transposons are inserted in the vicinity of endogenous gene loci. In the absence of Piwi or Panoramix, loss of repressive heterochromatin at these transposon insertions results in elevated transcription of these genes (Supplementary Fig. 2i; Table S4) (Sienski et al., 2012). A highly similar set of genes was differentially expressed in Nxf2- or Panoramix-depleted OSCs (Supplementary Fig. 2i; Table S4). The rare changes in gene expression caused by Nxf2 loss can, therefore, be attributed directly to impaired Piwi-mediated heterochromatin formation at transposon loci. We confirmed all findings based on RNAi-mediated depletion of Nxf2 in OSCs with a second independent siRNA targeting *nxf2* (Supplementary Fig. 2j, k; Table S4). Our combined results support a model where Nxf2—rather than acting as a cellular RNA transport receptor— is required for co-transcriptional silencing and heterochromatin formation downstream of Piwi.

### Targeting Nxf2 to nascent RNA induces co-transcriptional silencing

To test Nxf2’s involvement in heterochromatin-mediated silencing more directly, we converted a reporter system in fly ovaries that assays co-transcriptional silencing independently of piRNAs (Sienski et al., 2015; Yu et al., 2015) into a quantitative cell culture assay. We generated a clonal OSC line harboring a single-copy transgene that expresses GFP under control of a strong enhancer (Supplementary Fig. 3a). To mimic piRNA-guided target silencing, any factor of interest can be recruited to the nascent reporter RNA as a λN-fusion protein, through boxB sites located in the intron of the reporter construct (Fig. 3a). The same reporter cell line also allows recruitment of factors of interest to the reporter DNA (via the Gal4-UAS system) upstream of the enhancer in order to assay transcriptional silencing independent of targeting the nascent RNA (Fig. 3a).

**Figure 3.**
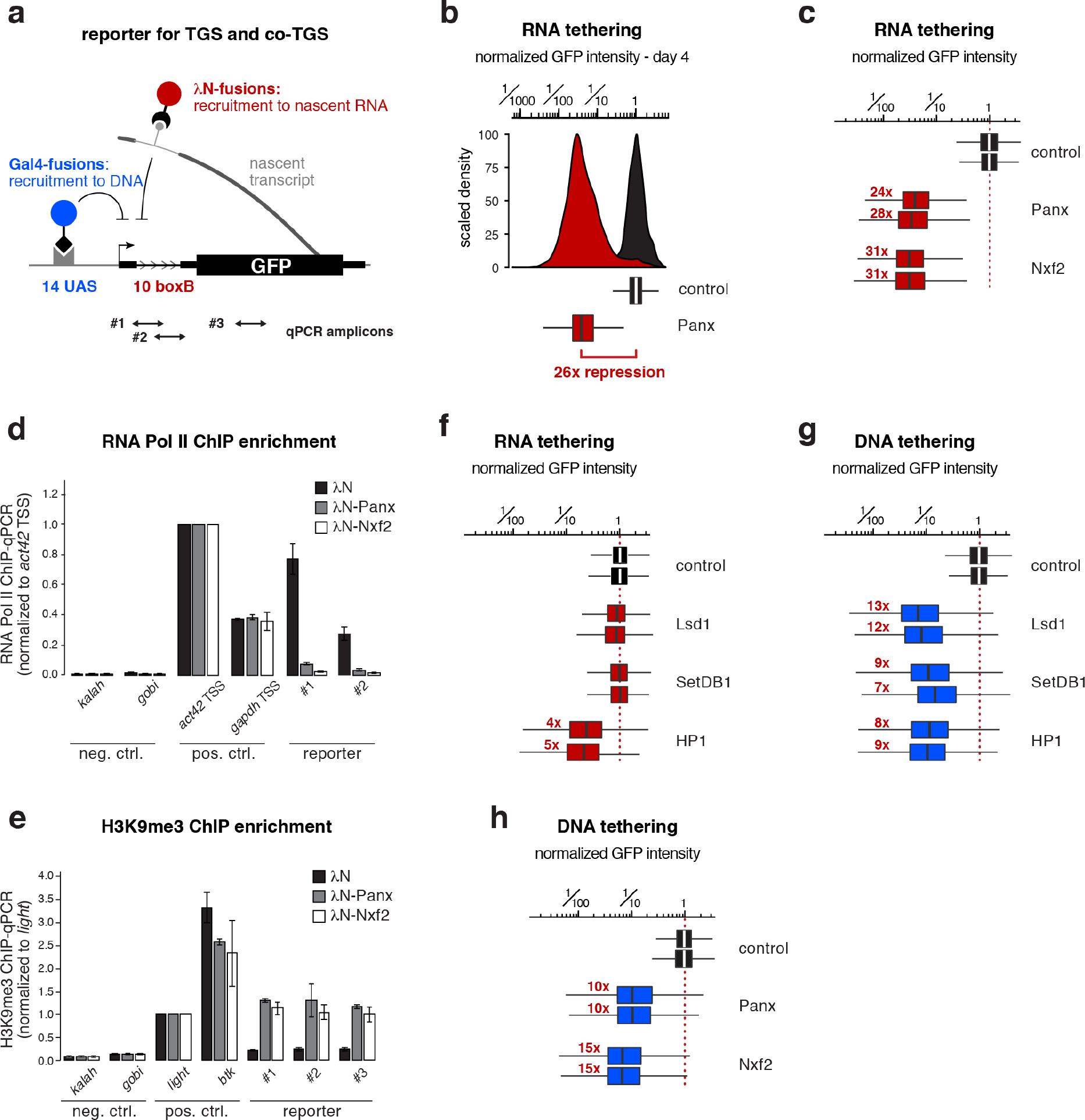
Targeting Nxf2 to nascent RNA induces co-transcriptional silencing. **a**, Cartoon of the GFP silencing reporter, which allows direct recruitment of proteins to the nascent RNA via intronic boxB sites and to DNA via UAS sites (positions of qPCR primer amplicons used in **d**, **e** are indicated). **b**, Histogram (top) and corresponding box plot (bottom) showing the λN-control normalized GFP intensity in transfected OSCs (4 days after transfection of λN-tagged fusion proteins; n=2500 cells; box plot definition as in Fig. 2b; red number indicates fold-repression value based on median, normalized to control). **c**, Box plots showing GFP intensity in OSCs 4 days after transfection with plasmids expressing indicated λN-fusion proteins (red numbers indicate fold-repression value based on median, normalized to control for two replicates; box plot definition as in Fig. 2b). **d, e**, Bar graphs showing normalized RNA Pol II occupancy (**d**) and H3K9me3 levels (**e**) at indicated loci determined by chromatin immunoprecipitation (ChIP) followed by quantitative PCR using OSCs expressing the indicated λN-fusions (n = 2; error bars: SD). **f**, As in c. **g, h**, As in c, but for indicated Gal4 DBD fusions.

While expression of λN alone had no impact on GFP reporter levels, transient expression of λN-Panoramix resulted in more than 25-fold reporter repression for four to five days (Fig. 3b; Supplementary Fig. 3b). Expression of λN-Nxf2 led to similarly potent repression (Fig. 3c; Supplementary Fig. 3c). In both cases, silencing correlated with reduced RNA Pol-II occupancy and establishment of an H3K9me3 domain at the reporter locus (Fig. 3d, e). Experimental tethering of either Panoramix or Nxf2 to a nascent RNA, therefore, induces co-transcriptional silencing accompanied by heterochromatin formation. The efficiency of this silencing process is remarkable considering that the boxB sites reside in an intron of the reporter construct, and thus one would expect that the target RNA is only transiently present at the encoding DNA locus. Indeed, the heterochromatin promoting factors Eggless/SetDB1, Su(var)3-3/Lsd1, or Su(var)205/HP1a, which act downstream of Piwi, were not capable of inducing comparable co-transcriptional silencing when targeted to the nascent reporter RNA (Fig. 3f). The same factors silenced the reporter as efficiently as Panoramix or Nxf2 when targeted directly to the reporter DNA (Fig. 3g, h). Therefore, we propose that co-transcriptional silencing, i.e. silencing via targeting nascent RNA, by Panoramix and Nxf2 requires more than merely recruiting the so-far identified heterochromatin effectors to the nascent RNA.

### Panoramix and Nxf2-Nxt1 form the interdependent SFiNX complex

Nxf2 and Panoramix interact, are both capable of inducing co-transcriptional silencing, and loss of either protein results in highly similar phenotypes (Fig. 1-3). These findings suggested a close molecular connection between Nxf2 and Panoramix. In support of this, we found a strong reciprocal dependency between both proteins: Depletion of Nxf2 or Panoramix in OSCs resulted in substantial reductions of the respective other protein, while the corresponding mRNA levels were unchanged (Fig. 4a, b). Similarly, in *nxf2* mutant ovaries, Panoramix protein was hardly detectable by western blotting or immunofluorescence analysis (Fig. 4c, d), despite unchanged *panoramix* mRNA levels (Supplementary Fig. 4a). Conversely, Nxf2 protein levels were reduced in *panoramix* mutant ovaries (Fig. 4c), and the remaining Nxf2 protein was excluded from the nucleus in germline and soma (Fig. 4e), suggesting that Nxf2’s nuclear localization depends on Panoramix. Consistent with this, Panoramix harbors a Lysine-rich sequence stretch (residues 196-262), which is required for its nuclear localization (Supplementary Fig. 4b). Based on these results, we hypothesized that Panoramix and Nxf2 stabilize each other via forming a nuclear protein complex.

**Figure 4.**
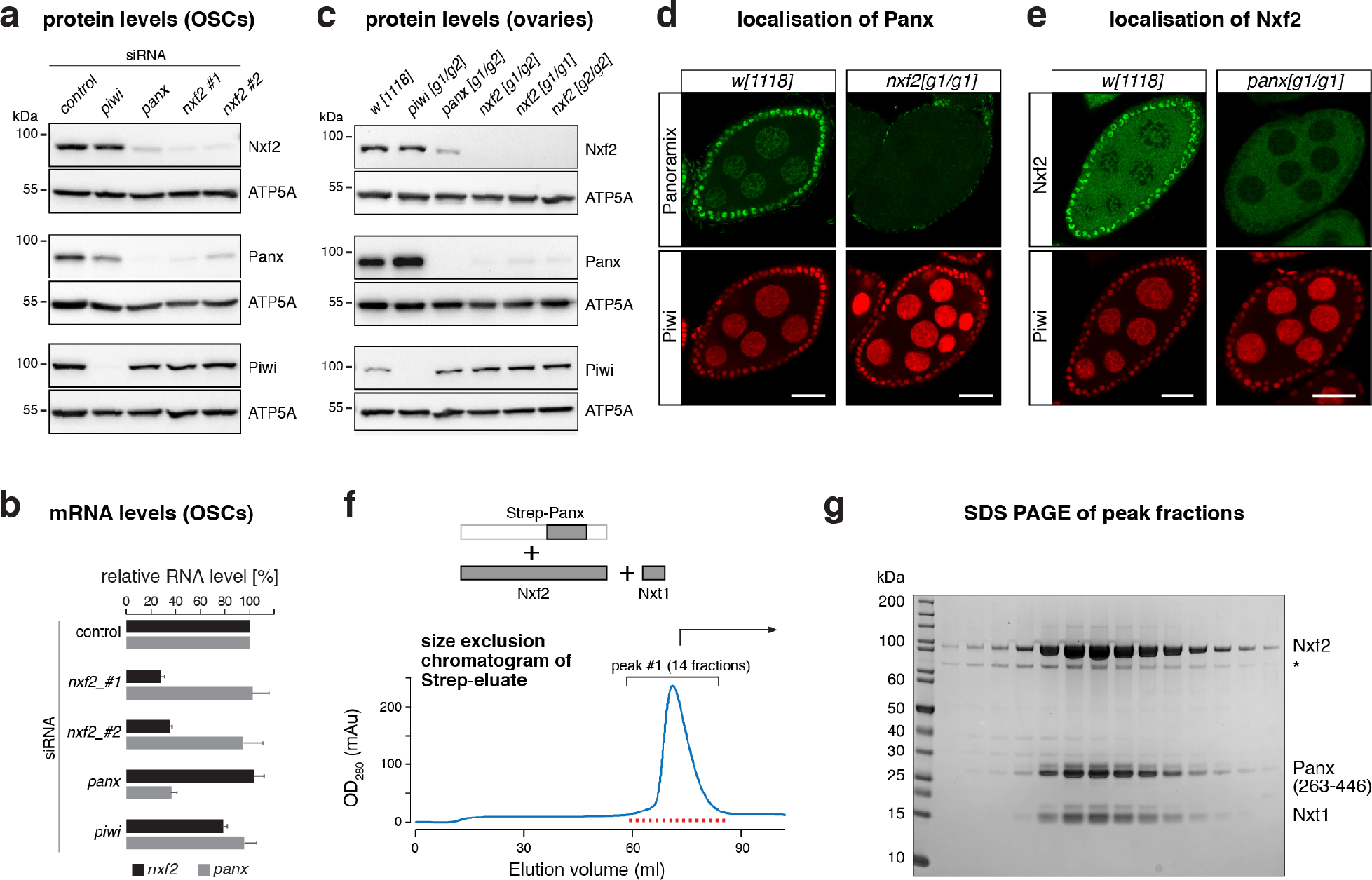
Panoramix and Nxf2-Nxt1 form the interdependent SFiNX complex. **a**, Western blots showing levels of indicated proteins in lysates from OSCs with indicated gene knockdowns (ATP5A served as loading control). **b**, Bar graph showing fold change of RNA-seq levels of indicated genes compared to control knockdown in OSCs (n = 3, error bars: SD). **c**, Western blots showing levels of indicated proteins in ovary lysates from flies with indicated genotypes (ATP5A served as loading control). **d**, Confocal images of egg chambers (scale bar: 20 μm) from *w[1118]* control and *nxf2* mutant flies stained for Panoramix (upper panels) and Piwi (lower panels). **e**, As in **d**, but for *panoramix* mutants and Nxf2 protein. **f**, Top: schematics of the trimeric complex expressed in insect cells (grey boxes indicate co-expressed proteins/fragments). Bottom: size exclusion chromatogram of the affinity-purified Strep-Panoramix eluate (mAU = milli-absorbance unit). **g**, SDS-PAGE of the 14 peak fractions indicated in **f** (Coomassie blue staining; * indicates HSP70 as determined by mass spectrometry).

To test for a putative Panoramix-Nxf2 complex, we co-expressed both proteins in insect cells together with Nxt1/p15, which we identified as Nxf2 and Panoramix interactor (Fig. 1), and which functions as a general NXF cofactor (Herold et al., 2000). While all three proteins were abundantly produced, Panoramix was degraded. Therefore, we expressed instead of full-length Panoramix a 25 kDa fragment that is necessary and sufficient for binding Nxf2 (see below). A single affinity purification step of the Strep-tagged Panoramix fragment, followed by size exclusion chromatography, resulted in a defined protein peak containing all three factors (Fig. 4f, g; Supplementary Fig. 4c). Panoramix, Nxf2, and Nxt1/p15 therefore form a stable protein complex, which we named SFiNX (<u>S</u>ilencing <u>F</u>actor <u>i</u>nteracting <u>N</u>uclear e<u>X</u>port factor variant).

### Panoramix-mediated silencing via nascent RNA requires Nxf2

Due to the interdependency between Panoramix and Nxf2, our experiments so far interrogated the function of the SFiNX complex rather than that of the individual proteins. To disentangle the molecular roles of Panoramix and Nxf2, we set out to generate interaction-deficient point-mutant variants. Panoramix consists of two parts, an N-terminal disordered half, and a C-terminal half with predicted secondary structure elements (Fig. 5a). Using GFP-tagged full-length Nxf2 as bait, we mapped the interaction site within Panoramix to the first part of the structured domain (Fig. 5a; Supplementary Fig. 5a, b). Within there, we identified two regions that upon deletion impacted the Nxf2-Panoramix interaction (Supplementary Fig. 5c). Panoramix Δ308-386 failed to bind Nxf2, while Panoramix Δ387-446 interacted less efficiently with Nxf2. The 308-386 peptide harbors a predicted amphipathic alpha-helix (Supplementary Fig. 5d, e). Mutating four hydrophobic residues predicted to line one side of this helix abrogated the Nxf2 interaction (Fig. 5b; Supplementary Fig. 5e). Notably, both Panoramix variants that are defective in Nxf2 binding (Panoramix[Δ308-386] and Panoramix[helix mutant]) accumulated to lower levels compared to the wildtype protein (Supplementary Fig. 5f). Panoramix Δ387-446 on the other hand accumulated to higher levels than wildtype Panoramix (Supplementary Fig. 5f). Considering the co-dependency between Panoramix and Nxf2 in vivo, we hypothesized that the 387-446 peptide within Panoramix harbors a destabilizing element that induces protein degradation if not protected by Nxf2. Consistent with this, fusing the 387-446 Panoramix peptide to GFP led to a ~100-fold reduction in GFP levels (Supplementary Fig. 5g). Mutation of four hydrophobic residues within this “degron” led to increased Panoramix levels (Supplementary Fig. 5d, f). When combining both sets of point mutations, the resulting Panoramix[helix+degron mutant] accumulated to high levels and was unable to interact with Nxf2 (Fig. 5b; Supplementary Fig. 5f).

**Figure 5.**
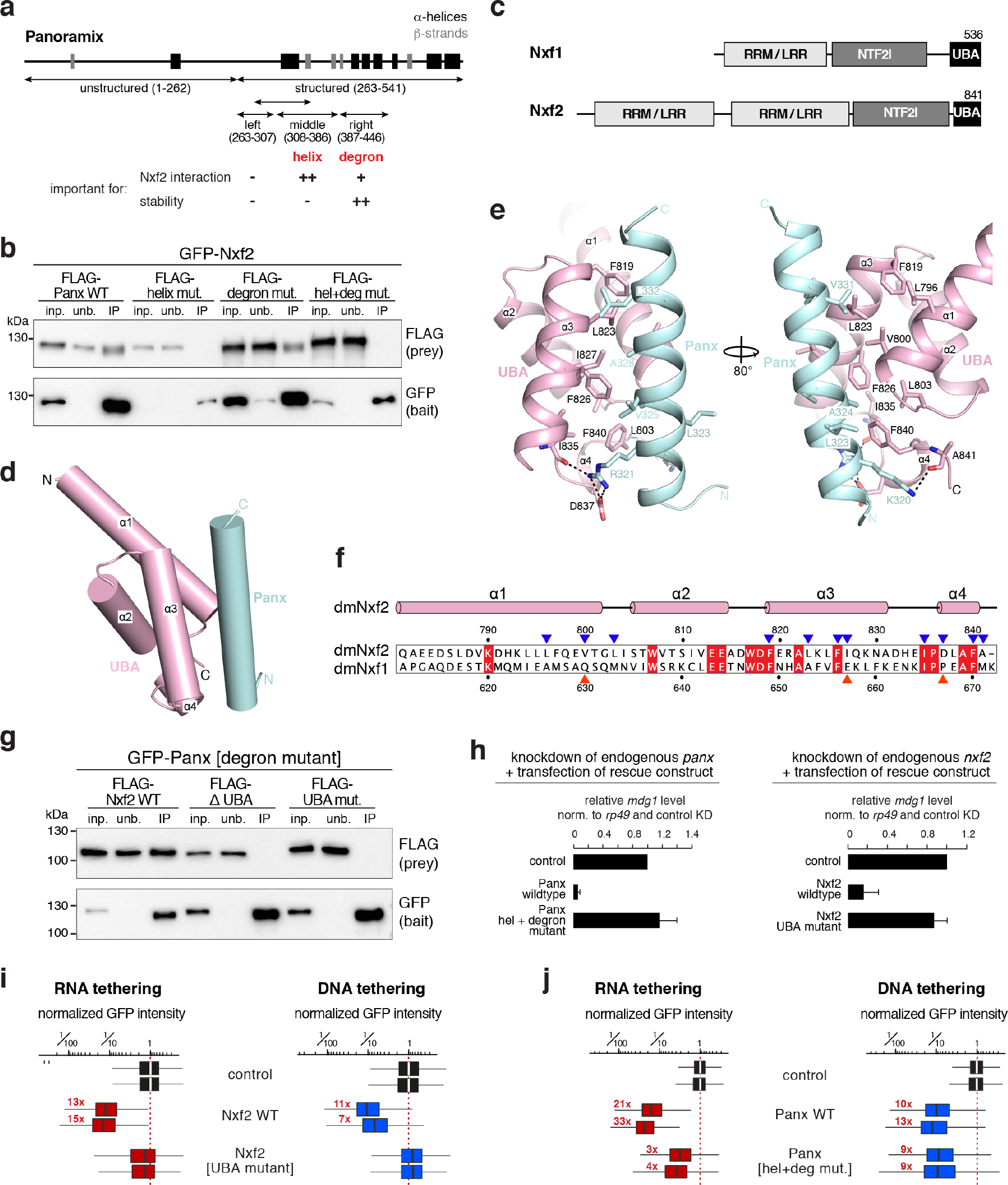
Panoramix-mediated silencing via nascent RNA requires Nxf2. **a**, Cartoon of the Panoramix protein showing predicted secondary structure elements and region boundaries used in the interaction mapping experiments. The summary below states the involvement of the [helix] and [degron] sites in the Nxf2 interaction and in Panoramix’ protein stability. **b**, Western blot analysis of GFP-Nxf2 immuno-precipitation experiments using lysate from S2 cells transiently co-transfected with indicated FLAG-Panoramix expressing plasmids (relative amount loaded in immunoprecipitation lanes: 3×). **c**, Schematics of the domain architecture of Nxf1/Tap and Nxf2. RRM: RNA recognition motif; LRR: leucine rich repeat; NTF2l: nuclear transport factor 2-like domain; UBA: ubiquitin associated domain. **d**, Cartoon view of the UBA domain of Nxf2 (in pink) in complex with the Panoramix helix (in cyan). **e**, Detailed views of interfaces between the UBA domain of Nxf2 (in pink) and the Panoramix helix (in cyan). **f**, Protein sequence alignment of the UBA domain of Nxf2 and Nxf1 from *Drosophila melanogaster* with the secondary structure elements of Nxf2 indicated above the alignment. Identical residues are highlighted by red squares, key Nxf2 residues involved in the Panoramix interaction are indicated by blue triangles, while red triangles mark interaction residues that are different in Nxf1. **g**, Western blot analysis of immuno-precipitation experiments (bait: stabilized degron mutant GFP-Panoramix) using lysate from S2 cells transiently co-transfected with indicated FLAG-Nxf2 expressing plasmids (relative amount loaded in immunoprecipitation lanes: 3×). **h**, Bar graphs showing the transposon repression rescue potential of indicated constructs transfected into OSCs depleted of endogenous Panoramix (left) or Nxf2 (right). *mdg1* transposon levels determined via qRT-PCR (n = 2, error bars: SD). **i, j**, Box plots showing GFP intensity in OSCs 4 days after transfection with plasmids expressing indicated λN-tagged (red) or Gal4-tagged (blue) fusion proteins (red numbers indicate fold-repression values based on median, normalized to control for two replicates; box plot definition as in Fig. 2b).

We then turned to Nxf2, whose domain organization resembles that of the mRNA export receptor Nxf1/Tap, except that it has a duplicated putative RNA binding unit consisting of RNA recognition motif (RRM) and a Leucine-rich repeat (LRR) domain (Fig. 5c). Using stabilized Panoramix[degron-mutant] as bait, we determined that Nxf2’s NTF2-like and UBA domains together are sufficient to bind Panoramix, and that the UBA domain is required for the Panoramix interaction (Supplementary Fig. 5h). Consistent with this, the 30 amino-acid amphipathic Panoramix helix, that is required for the Nxf2 interaction (Fig. 5b), bound the Nxf2 UBA domain in vitro (Supplementary Fig. 5i). Building on this, we determined the structure of Nxf2’s UBA domain in complex with the Panoramix helix at 2.0 Å resolution (Supplementary Fig. 6a, b; Table S5). This revealed that the UBA domain of *Drosophila* Nxf2 is highly similar to that of human NXF1 (PDB ID: 1OAI; Grant et al., 2003). Both domains consist of a three-helix bundle (α1-α3) with a short fourth helix (α4) at the C terminus (Fig. 5d). α1, α3 and α4 of the Nxf2 UBA domain form a hydrophobic core that interacts with the hydrophobic face of the amphipathic Panoramix helix involving the Panoramix residues L323, A324, V325, A328, V331, and L332. In addition, the N terminus of the Panoramix helix is stabilized by two flanking hydrogen bonds and salt bridges (Fig. 5e). The experimentally determined Panoramix[helix mutant] variant that cannot bind Nxf2 (Fig. 5b) fully supported this structure: three out of the six hydrophobic residues contributing to the interaction were mutated in the Panoramix[helix mutant] variant. Within Nxf2, ten hydrophobic residues contribute to the Panoramix interaction (Fig. 5f). Out of these, two residues (V800 and I827) are highly different in *Drosophila* Nxf1 (hydrophilic residues Q632 and E657), which does not interact with Panoramix (Supplementary Fig. 6c, d). When we mutated V800 and I827 in Nxf2, together with two flanking residues, into the corresponding Nxf1 amino acids, the resulting Nxf2[UBA mutant] was unable to bind Panoramix (Fig. 5g; Supplementary Fig. 6c).

Building on the interaction-deficient Nxf2 and Panoramix variants, we determined whether the individual proteins are capable of supporting Piwi-mediated silencing. We performed genetic rescue experiments in OSCs and asked whether expression of siRNA resistant Panoramix[helix+degron-mutant] or Nxf2[UBA-mutant] variants could restore silencing of the *mdg1* transposon in OSCs depleted for endogenous Panoramix or Nxf2. While the respective wild-type proteins supported *mdg1* silencing, neither of the interaction-deficient mutants (expressed with NLS sequence to assure nuclear localization; Supplementary Fig. 6e, f) displayed rescue activity (Fig. 5h). Therefore, both Nxf2 and Panoramix contribute essential activities to the silencing process beyond reciprocal protein stabilization. To investigate the function of Panoramix and Nxf2 as silencing factors more directly, we turned to the transcriptional silencing reporter assay in OSCs (Fig. 3a). Nxf2 with point mutated UBA domain was entirely inert in inducing reporter silencing, irrespective of whether it was targeted to the nascent RNA or to the DNA directly (Fig. 5i; Supplementary Fig. 6g). Instead, the Panoramix [helix+degron mutant] variant, which is defective in Nxf2 binding, showed clear, though in comparison to the wildtype protein only weak co-transcriptional silencing activity (Fig. 5j; Supplementary Fig. 6h). Remarkably, when recruited directly to the reporter DNA, Panoramix[helix+degron mutant] was as potent in inducing silencing as wildtype Panoramix (Fig. 5j; Supplementary Fig. 6h). Taken together, our data indicate that Panoramix, and not Nxf2, connect SFiNX to the downstream silencing machinery. Nxf2 instead is required for Panoramix to achieve potent silencing via the nascent RNA. To understand the function of Nxf2 within SFiNX, we reasoned that a specific molecular feature intrinsic to Nxf1/Tap was exploited by the evolutionary exaptation of an RNA transporter into co-transcriptional silencing. At the same time, Nxf2 must have lost other Nxf1 characteristics in order to not get channeled into mRNA export biology. We therefore set out to systematically compare Nxf2 to the well-studied Nxf1/Tap protein.

### The Nxf2-Nxt1 heterodimer lost nucleoporin binding activity

A central molecular feature of Nxf1/Tap is its ability to shuttle through the selective phenylalanine-glycine (FG) repeat meshwork of the inner nuclear pore complex (NPC). Two nucleoporin FG-binding pockets, one residing in the UBA domain and one in the NTF2-like domain, confer NPC shuttling ability to Nxf1/Tap (Braun et al., 2002). We examined both sites in Nxf2 at the structural level. The putative FG-binding pocket within Nxf2’s UBA domain lies on the opposite side of the Panoramix binding surface, making it per se accessible (Fig. 6a). However, a salt bridge between E814 and K829 restricts access to the hydrophobic core of the pocket, rendering it most likely non-functional (Fig. 6b). To inspect the second putative FG-binding pocket, we determined the 2.8 Å resolution crystal structure of Nxf2’s NTF2-like domain bound to Nxt1/p15 (Supplementary Fig. 7a; Table S5). Based on this, Nxf2 interacts with Nxt1/p15 in a manner very similar to human NXF1/TAP (Fig. 6c) (Fribourg et al., 2001). But in contrast to the human protein, Nxf2’s putative FG-binding pocket within the NTF2-like domain is again concealed: It is occupied by the bulky side chains of its own Phe735 and Tyr690 and in addition, Arg747 closes access to the hydrophobic pocket by hydrogen bonding with Tyr690 (Fig. 6d). As mutations in either of the two FG-binding pockets abrogate nucleoporin binding for human NXF1/TAP (Braun et al., 2002), it is highly unlikely that *Drosophila* Nxf2 retained NPC binding ability. Indeed, in vitro experiments demonstrated lack of affinity between FG-repeat peptides and Nxf2 (in complex with Nxt1 and Panoramix), while human NXF1/TAP (in complex with NXT1) readily bound FG-peptides (Fig. 6e). Furthermore, GFP-tagged Nxf2 did not accumulate at nuclear pores where GFP-tagged Nxf1/Tap is highly enriched (Supplementary Fig. 7b). Taken together, our biochemical and structural data indicate that Nxf2 lost nucleoporin binding activity, an evolutionary change that is in line with the repurposing of this NXF variant for co-transcriptional silencing.

**Figure 6.**
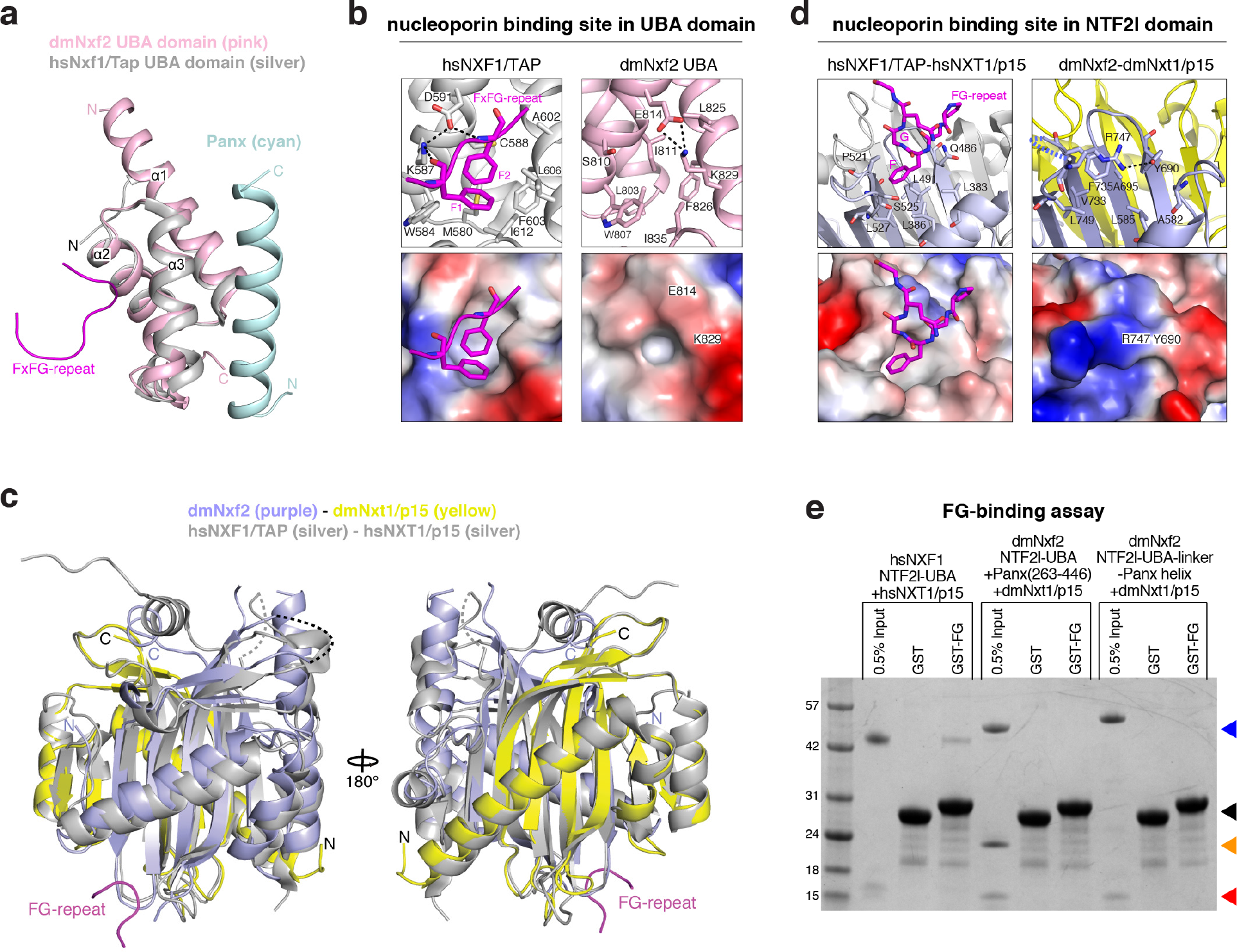
Nxf2 lost nucleoporin binding. **a**, Superposition of the overall structure of dmNxf2 UBA domain (in pink) complexed with Panoramix helix (in cyan) and the crystal structure of hsNXF1/TAP UBA domain (PDB ID: 1OAI) (in silver) (Grant et al., 2003). **b**, Left: Ribbon (top) and electrostatic surface (bottom) representation showing the FG-repeat binding pocket in the crystal structure of hsNXF1/TAP UBA domain (in gray) in complex with a nucleoporin FxFG-repeat peptide (magenta; PDB ID: 1OAI) (Grant et al., 2003). Right: Ribbon (top) and electrostatic surface (bottom) representation showing the closed putative FG-repeat binding pocket in the crystal structure of dmNxf2’s UBA domain (in pink). **c**, Superposition of the overall structure of dmNxf2’s NTF2-like domain complexed with dmNxt1/p15 (in purple and yellow) and the crystal structure (in silver) of hsNXF1/TAP-NXT1/p15 (PDB ID: 1JKG) (Fribourg et al., 2001). Invisible loops in the structure are shown as dashed curves. **d**, Left: Ribbon (top) and electrostatic surface (bottom) representation showing the specific recognition of nucleoporin FG-repeat (magenta) in the hsNXF1/TAP (in light blue)-NXT1/p15 (in gray) structure (PDB ID: 1JN5) by a hydrophobic FG-repeat binding pocket (Fribourg et al., 2001). Right: Ribbon (top) and electrostatic surface (bottom) representation showing the blocked putative FG-repeat binding pocket in the crystal structure of Nxf2’s NTF2-like domain (in purple) complexed with Nxt1/p15 (in yellow). The hydrogen bond is shown as black dashed line. **e**, Coomassie-stained SDS PAGE testing the interaction between FG-peptides and indicated protein complexes (arrowheads at the right refer to: black: GST or GST-FG repeat; blue: NXF1/TAP, Nxf2(NTF2l + UBA) or Nxf2(NTF2l + UBA)-linker-Panoramix-helix); red: Nxt1; orange: Panoramix (263-446).

### A key role for Nxf2’s RNA binding unit within SFiNX

Nxf1/Tap’s second core feature is its ability to bind the mRNA cargo via the N-terminal RRM-LRR domains (Liker et al., 2000). RNA binding is non-sequence-specific and requires recruitment of Nxf1/Tap to cargo RNA through various adaptor proteins that mark the completion of the different mRNA processing steps. Nxf2 harbors a tandem RRM-LRR fold (Fig. 5c). Based on electrophoretic mobility shift assays, Nxf2’s N-terminal RRM-LRR domain (1^st^ unit) is capable of binding single-stranded RNA in vitro (Fig. 7a, b). The RNA binding activity of the 1^st^ RRM-LRR unit was abrogated upon mutating three positively charged amino acids, whose equivalent residues in NXF1/TAP contact the constitutive transport element (CTE) of simian type D retroviral transcripts (Supplementary Fig. 8a-c) (Aibara et al., 2015; Teplova et al., 2011). As we did not succeed in obtaining Nxf2’s second RRM-LRR unit as a soluble recombinant protein, it is currently unclear whether this unit provides additional RNA binding activity.

**Figure 7.**
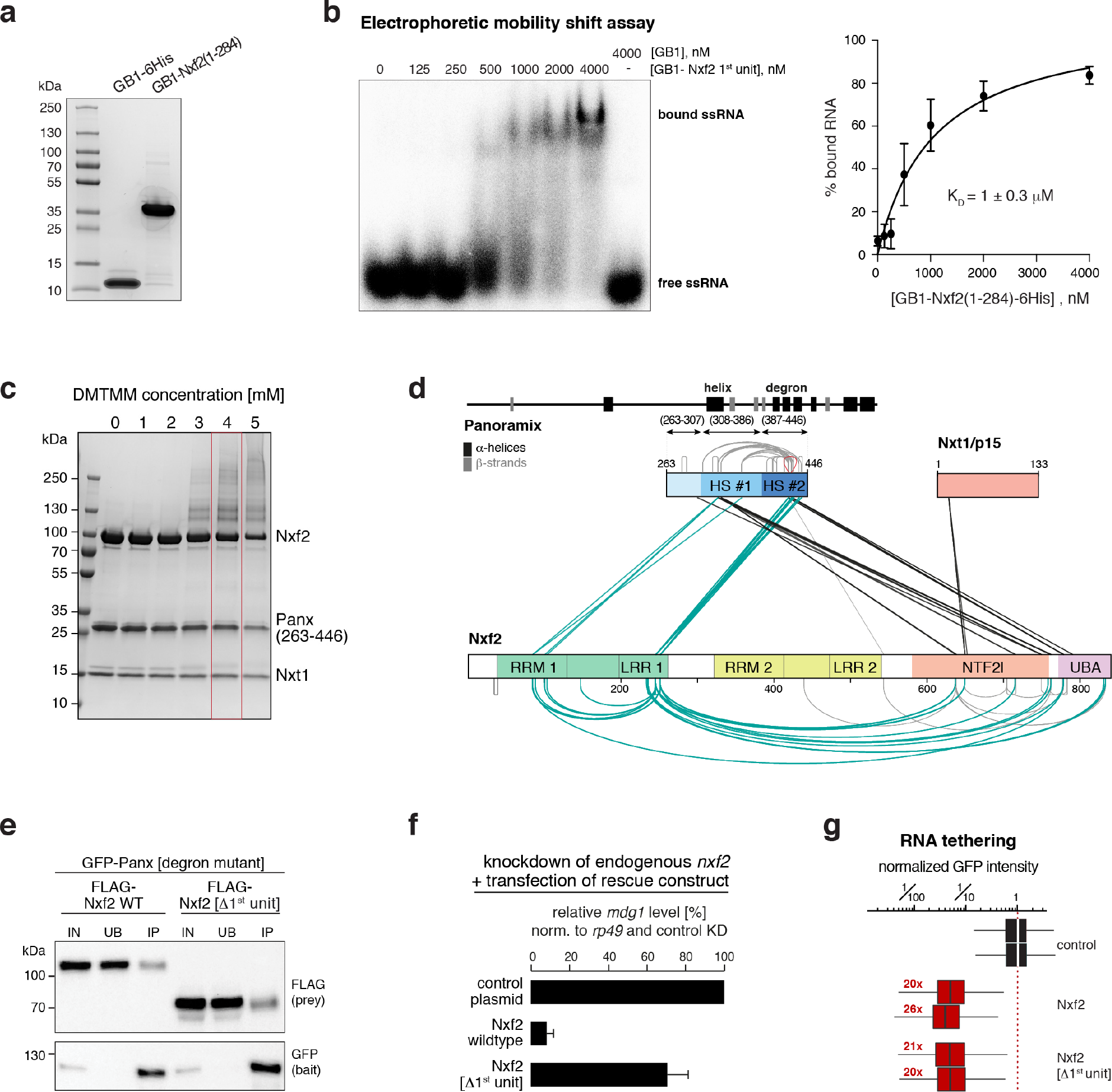
A key role for Nxf2’s RNA binding unit within SFiNX. **a**, Coomassie stained SDS-PAGE showing purified RNA-binding unit (1^st^ RRM-LRR domain) of Nxf2 and the GB1 control peptide used for the EMSA experiment (b). **b**, Left: Phosphorimage showing Electrophoretic Mobility Shift Assay (EMSA) with labelled single stranded RNA (ssRNA) and increasing amount of indicated recombinant protein. Right: Binding curve for the EMSA calculated from the phosphorimage (n = 3; error bars: SD). **c**, SDS-PAGE (Coomassie blue staining) showing chemically crosslinked recombinant SFiNX complex. The condition boxed in red was chosen for the crosslinking and mass spectrometric analysis. **d**, Schematic summary of all crosslinks identified between Panoramix, Nxf2, and Nxt1 in the recombinant SFiNX complex. Protein domains in Nxf2 are indicated (RRM: RNA recognition motif; LRR: leucine rich repeat; NTF2l: nuclear transport factor 2-like domain; UBA: ubiquitin associated domain). Within Panoramix, hotspot (HS) #1 includes the amphipathic helix, and HS #2 includes the degron site. Numbers indicate amino acid positions within the proteins. Straight lines indicate intermolecular crosslinks, half-circles indicate intramolecular crosslinks and the red loop indicates an intermolecular self-link. **e**, Western blot analysis of immuno-precipitation experiments (bait: stabilized degron-mutant GFP-Panoramix) using lysate from S2 cells transiently co-transfected with indicated FLAG-Nxf2 expressing plasmids (relative amount loaded in immunoprecipitation lanes: 3x). **f**, Bar graph showing the transposon repression rescue potential of indicated constructs transfected into OSCs depleted of endogenous Nxf2. *mdg1* transposon levels determined via qRT-PCR (n = 2, error bars: SD). **g**, Box plots showing GFP intensity in OSCs 4 days after transfection with plasmids expressing indicated λN-tagged fusion proteins (red numbers: fold-repression values based on median, normalized to control for two replicates are indicated; box plot definition as in Fig. 2b).

Nxf2’s ability to bind RNA raised the important question of how it avoids binding to random nuclear RNAs, which would bear the danger of ectopic silencing and heterochromatin formation. Inspired by the Nxf1 literature, we hypothesized that Nxf2’s RNA binding activity is regulated. It has been suggested that prior to mRNA cargo binding, Nxf1/Tap is in a closed conformation with its RNA binding unit folding back onto the NTF2-like domain (Viphakone et al., 2012). Upon adaptor-mediated recruitment of Nxf1/Tap to an export-competent mRNA, this intramolecular inhibition is released, RNA cargo is bound, and the complex shuttles through the NPC. To probe for a putative intramolecular regulatory interaction within SFiNX, we took advantage of the recombinant Panoramix-Nxf2-Nxt1 complex (Fig. 4). We used chemical crosslinking coupled to mass spectrometry in order to establish an interaction map of residues that are in close physical proximity within the complex (Fig. 7c, d; Table S6). One set of identified crosslinks (black in Fig. 7d) was in full agreement with our structural and biochemical data. First, the two crosslinks involving Nxt1 map to the NTF2-like domain. Second, two crosslinking hotspots are apparent within Panoramix. Hotspot #1 corresponds to the amphipathic helix, hotspot #2 to the degron site. Both hotspots exhibit several crosslinks to the C-terminus of Nxf2, indicating that the Panoramix-Nxf2 interaction involves besides the helix-UBA interaction that we characterized (Fig. 5), additional interactions, probably with the NTF2-like domain. Strikingly, nearly all other identified protein crosslinks involve Nxf2’s first RRM-LRR unit, which according to our in vitro experiments is also capable of RNA binding. We identified multiple intramolecular crosslinks between the first RRM-LRR unit and the NTF2-like and UBA domains, as well as intermolecular crosslinks to Panoramix, mostly to the degron site. As the 1^st^ RRM-LRR unit is dispensable for Panoramix binding (Fig. 7e), the identified intra-molecular interactions point to a regulatory interaction, similar to what is proposed for Nxf1/Tap (Viphakone et al., 2012).

To determine the importance of the first RRM-LRR unit for SFiNX function, we performed genetic rescue experiments. In flies and OSCs, expression of Nxf2[Δ1^st^ RRM-LRR] instead of the wildtype protein was not able to support transposon silencing (Fig. 7f; Supplementary Fig. 8d, e), although Nxf2[Δ1^st^ RRM-LRR] localized to the nucleus (Supplementary Fig. 8d) and interacted with Panoramix (Fig. 7e). Together with the finding that the first RRM-LRR unit is able to bind RNA, this suggested a model where Nxf2 anchors SFiNX to the nascent target RNA via its N-terminal RRM-LRR domain. If this were true, SFiNX lacking Nxf2’s 1^st^ RNA binding unit should remain silencing competent if recruited to the target RNA via the λN-boxB tethering system. Indeed, expression of λN-tagged Nxf2[Δ1^st^ RRM-LRR] induced co-transcriptional silencing as efficiently as λN-tagged wildtype Nxf2 (Fig. 7g; Supplementary Fig. 8f). Taken together, our findings are consistent with a model where in the non-target-engaged state, Nxf2’s N-terminal RNA binding unit is in an auto-inhibited state by folding back onto the SFiNX complex. Upon recruitment to a target transcript, Nxf2’s N-terminal RRM-LRR domain would interact with RNA, thereby anchoring SFiNX via the nascent target transcript to chromatin and allowing Panoramix to recruit effectors to establish heterochromatin (Supplementary Fig. 8g).

## DISCUSSION

The discovery of SFiNX, a nuclear protein complex consisting of Panoramix, Nxf2, and Nxt1/p15, provides the key molecular connection between Piwi, the nascent target RNA, and the cellular heterochromatin machinery. In the absence of SFiNX, piRNA-loaded Piwi is incapable of inducing co-transcriptional silencing (Fig. 2). Conversely, experimental recruitment of SFiNX, but not its individual components, to a nascent RNA results in potent silencing and local heterochromatin formation independently of Piwi (Fig. 3). Our data indicate that within SFiNX, Panoramix, and not Nxf2, provides the molecular link to the downstream cellular heterochromatin effectors (Fig. 5). Based on genetic experiments, the histone methyl-transferase SetDB1/Eggless, the histone-demethylase Lsd1/Su(var)3-3, and the heterochromatin binding protein HP1/Su(var)205 are required for piRNA-guided co-transcriptional silencing (Sienski et al., 2015; Yu et al., 2015). We did not find any of these factors enriched in our SFiNX co-IP mass-spectrometry experiments. Interestingly, the SUMO E3 ligase Su(var)2-10, which was recently linked to piRNA-guided co-transcriptional silencing (Ninova et al., 2019), is enriched more than five-fold in Nxf2 and Panoramix IP experiments (Table S1).

The involvement of Nxf2, a nuclear RNA export variant, in co-transcriptional silencing came as a surprise to us. Based on our biochemical and structural data, we propose that two molecular features of the ancestral NXF protein, the principal mRNA export receptor Nxf1/Tap, facilitated the evolutionary exaptation of Nxf2 into piRNA-guided silencing (Fig. 7). First, Nxf2 retained its ability to bind RNA, thereby providing SFiNX a molecular link to the nascent target RNA. Consistent with this, Siomi and colleagues provide evidence based on CLIP experiments that Nxf2 directly interacts with piRNA complementary target RNAs in OSCs (Murano et al., 2019). Second, our crosslinking-mass spectrometry data indicate that similar to Nxf1/Tap, the RNA binding activity of Nxf2 might be gated. We propose that this could provide a critical regulatory switch to ensure that SFiNX only associates with transcripts that are specified as targets via the Piwi-piRNA complex. A central open question is, how SFiNX is recruited to piRNA complementary target RNAs, and whether the RRM-LRR units are involved in this recruitment. In the case of Nxf1/Tap, various proteins (e.g. SR-proteins, THO-complex, UAP56, Aly/Ref) that are recruited during co-transcriptional mRNA maturation, are required to restrict Nxf1/Tap deposition onto export-competent mRNAs only (Cullen, 2000; Heath et al., 2016; Kohler and Hurt, 2007). Whether any of these factors is also required for loading Nxf2 onto RNA is currently unclear. It is, however, likely that target-engaged Piwi contributes a critical role in the deposition of SFiNX onto the target RNA, potentially by licensing Nxf2’s RNA binding activity. We note that despite considerable experimental efforts we were not able to establish a direct molecular link between Piwi and either Nxf2 or Panoramix, although small amounts of Piwi consistently co-immuno-precipitate with SFiNX. Considering that Piwi represses transcription of its targets, the steady state level of a Piwi-SFiNX complex is expected to be small. In support of this, piRNA-independent recruitment of Piwi to a nascent RNA is incapable of target silencing, indicating that the Piwi-SFiNX interaction occurs only once Piwi is bound to a target RNA via a complementary piRNA (Post et al., 2014; Sienski et al., 2015; Yu et al., 2015). Further biochemical experiments, as well as structural insight into the full SFiNX complex will be required to shed light onto the molecular logic of this intriguing silencing complex.

Although no direct ortholog of Nxf2 is identifiable in vertebrates, our finding that an NXF variant is involved in transposon silencing in *Drosophila* likely points to a more general scheme. The Nxf1/Tap ancestor diversified through several independent evolutionary radiations into numerous NXF variants in different animal lineages (Supplementary Fig. 8h; Table S7). In flies, the three NXF variants exhibit gonad-specific expression with Nxf2 and Nxf3 being expressed predominantly in ovaries, and Nxf4 being testis-specific (Gramates et al., 2017). Besides Nxf2 (this study), *Drosophila* Nxf3 is also an essential piRNA pathway component as it is required for the nuclear export of un-processed piRNA cluster transcripts in the germline (ElMaghraby et al., 2019). In mice and humans, several NXF variants are preferentially expressed in testes. Intriguingly, Nxf2 mutant mice are male sterile, a phenotype shared with many piRNA pathway mutants (Pan et al., 2009). Considering this, we speculate that also in vertebrates the host-transposon conflict has been a key driver of NXF protein family evolution through frequent duplication and exaptation events. Our study highlights that some of these variants might have evolved novel molecular functions, not directly related to RNA export biology.

## ACKNOWLEDGMENTS

We thank K. Meixner for experimental support, P. Duchek and J. Gokcezade for generating CRISPR edited and transgenic flies, the VBCF NGS unit for deep sequencing, VBCF Protein Technologies Facility for recombinant protein expression, the MFPL monoclonal facility for Nxf2 and Panoramix antibodies, and the VDRC, TRiP, and Bloomington stock centers for flies. We thank A. Koehler and G. Riddihough from Life Science Editors (http://lifescienceeditors.com) for comments on the manuscript. We thank the Brennecke lab, particularly P. Andersen, for support and feedback. The Brennecke lab is supported by the Austrian Academy of Sciences, the European Community (ERC-2015-CoG - 682181), and the Austrian Science Fund (F 4303 and W1207). J. Batki was supported by the Boehringer Ingelheim Fonds. X-ray diffraction studies were conducted at the Advanced Photon Source on the Northeastern Collaborative Access Team beamlines, which are supported by NIGMS grant P30 GM124165 and U.S. Department of Energy grant DE-AC02-06CH11357. The Pilatus 6M detector on 24-ID-C beam line is funded by a NIH-ORIP HEI grant (S10 RR029205). MSKCC core facilities are supported by P30 CA008748. This work was supported by funds from the Maloris Foundation (DJP) and MSKCC core grant (P30 CA008748).

## AUTHOR CONTRIBUTIONS

J.Batki, J.S., A.I.V, L.L., and K.K. performed all molecular biology and fly experiments. D.H., J.S., and J.Batki performed the computational analyses, C.S. and K.M. performed the X-link mass spectrometry analysis, M.N. generated the phylogenetic comparisons of NXF proteins. J.W. generated, purified and grew crystals of the UBA-linker-helix and the Nxf2 NTF2l-Nxt1 complex, performed the X-ray crystallographic analyses, the GST pull-down experiments and performed the SEC-MALS assay with W.X., under the supervision of D.J.P. The paper was written by J.Batki, J.S. and J.B. with input from J.W. and D.J.P.

## MATERIALS & METHODS

### Fly strains

All fly strains used in this study are listed in Table S8. Flies were kept at 25 °C. For each experiment, flies were aged for 5-6 days and kept on apple juice agar plates with yeast paste to ensure consistent ovarian morphology. Two independent *nxf2* frameshift mutant alleles were generated by injecting the pDCC6 plasmid with *nxf2* targeting gRNAs (Table S9) into *w[1118]* flies (Gokcezade et al., 2014). Sequences for the two frameshift alleles are indicated in Supplementary Fig. 1c. N-terminal 3xFLAG_V5_GFP-tagging of endogenous *nxf2* and *panoramix* loci was done by co-injecting a repair template and the gRNA containing plasmid (Addgene 45956) into act-Cas9 flies (BL-58492). Oligonucleotides used for gRNA cloning are listed in Table S9.

Germline and soma specific gene knockdowns were performed by crossing short hairpin (shRNA) transgene strains with the maternal triple driver (MTD)-GAL4 line or the *traffic jam*-GAL4 driver line, respectively. Oligonucleotides used for shRNA cloning are listed in Table S9. *gypsy* and *burdock*-LacZ sensor strains are described in (Handler et al., 2013).

Rescue strains with different Panoramix and Nxf2 variants were generated by injecting respective rescue transgenes into *panoramix* or *nxf2* mutant flies containing attp landing sites on the same chromosome (attP40 for *panoramix*, attP154 for *nxf2*). Rescue transgenes contained the *panoramix* regulatory control regions (chr2R: 21,308,437-21,313,490) and the *panoramix* coding sequence was replaced by the various *panoramix* and *nxf2* variants.

### X-gal staining of ovaries

Ovaries were dissected into ice cold PBS, fixed in 0.5% glutaraldehyde (in PBS) for 15 min at room temperature, and then washed twice with PBS. Next, samples were incubated in staining solution (10 mM PBS, 1 mM MgCl2, 150 mM NaCl, 3 mM potassium ferricyanide, 3 mM potassium ferrocyanide, 0.1% Triton X-100, 0.1% X-gal (5-bromo-4-chloro-3-indolyl-β-d-galactoside)) overnight (*gypsy*-sensor) or for 2h (*burdock*-sensor) at room temperature.

### Generation of Nxf2 and Panoramix antibodies

Purified His-tagged Nxf2 (1-326) protein was used to generate the mouse anti-Nxf2 antibody used for western blot. The mouse anti-Nxf2 and anti-Panoramix antibodies used for immunofluorescence were raised against the SFiNX complex consisting of his-tagged Nxf2 (541-841), Strep-tagged-Panoramix(263-446) and Flag-tagged Nxt1 (full length). All antibodies were generated at the MFPL Monoclonal Antibody Facility.

### OSC cell culture

OSCs were cultured as described (Niki et al., 2006; Saito et al., 2009). Plasmid and siRNA transfections were performed using Cell Line Nucleofector kit V (Amaxa Biosystems) with the program T-029, using 8 million cells per transfection. siRNAs used in this study are listed in Table S10.

### Stable OSC reporter line generation

The reporter construct (*traffic jam* enhancer driven GFP_P2A-Blasticidin-resistance harboring 10 intronic boxB sites and 14 upstream UAS sites; plasmid submitted to Addgene) was integrated into chromosomal location chr2L:9,094,918, which is devoid of genes and major chromatin marks, using CRISPR-Cas9. In brief, 600bp long homology arms flanking the integration site were amplified from OSC genomic DNA. Oligonucleotides targeting the locus (Table S9) were cloned into the guide RNA expression plasmid (Addgene 49330). Two independent gRNA containing plasmids were mixed 1:1 and 200 ng of this mix were co-transfected with 1200 ng of the integration plasmid into OSCs. After two days, the cells were plated with different dilutions and on the following day Blasticidin containing media was added (1:1000) for a 4-day long selection. Afterwards, the cells were grown in normal medium for about 2 weeks until individual clones could be isolated.

### Droplet PCR

To assess the copy number of the reporter construct integrated in the OSC genome, the QX200™ Droplet Digital™ PCR System (BIORAD) was used according to the manufacturer’s instructions. In brief, genomic OSC DNA was digested with EcoRI and HindIII restriction enzymes. The PCR reaction was set up with 10ng digested genomic DNA (primer sequences in Table S9) and the QX200™ ddPCR™ EvaGreen Supermix. The PCR mix and the QX200 Droplet Generation Oil for EvaGreen were added into a DG8™ cartridge and droplets were generated with QX200 Droplet generator. Thermal cycling and droplet reading were performed with the instructor’s standard protocol which gave the concentration of the amplicon in copies/reaction volume. Based on 2 house-keeping control genes, the copy number in the genome was determined for the integrated reporter.

### Stable OSC line generation with extra genomic copy

The pAcm vector (Saito et al., 2009) was modified to create the integration constructs. Downstream of the *act5C* promoter, a 3xFLAG-HA tag was added followed by the open reading frames encoding full length Panoramix or Nxf2. The selection cassette consisted of an independent transcription unit driving mCherry_P2A_Puromycin-resistance via the *traffic jam* enhancer from *Drosophila yakuba* in combination with the *Drosophila* synthetic core promoter (DSCP) (Pfeiffer et al., 2008). The two transgenes were integrated into the chromosomal location chr2L:9,103,945, which is devoid of genes. 1200 bp long homology arms flanking the integration sequence were used for the integration. Stable integration was generated as described above for the reporter cell line, except that Puromycin was used for the selection (1:2000).

### Tethering reporter assay

All tethering constructs are based on λN-entry or Gal4-entry vectors (submitted to Addgene). Various full-length CDS or CDS variants were inserted into the entry vectors to generate N-terminally tagged fusion proteins. Unless having full length genes, the SV40 NLS sequence (PKKKRKV) was included to ensure nuclear localization of the variants. A separate expression cassette driving mCherry via the *traffic jam* enhancer was used to select positively transfected cells. OSCs harboring stably integrated GFP reporter were transfected with 4µg of λN/Gal4 fusion construct. As a negative control, λN/Gal4 empty vector was used. Two days after transfection, cells were harvested for WB analysis and four days after transfection cells were harvested for flow cytometry analysis using a FACS BD LSR Fortessa (BD Biosciences). Transfected cells were gated based on mCherry expression and the GFP intensity was determined in that population (per experiment 2500 cells). Data analysis was performed using FACS Diva and FlowJo.

### OSC rescue assay

OSCs were co-transfected with siRNAs targeting *panoramix* or *nxf2* and a plasmid containing the *act5c* driven siRNA-resistant rescue construct. A second transfection was performed after two days and cells were collected after four days for WB analysis and RNA isolation for RT-qPCR. siRNAs used for the rescue experiments are listed in Table S10.

### RT-qPCR

OSCs or 5-10 pairs of ovaries were collected into TRIzol reagent and RNA was isolated according to the manufacturer’s instructions. Total RNA was digested with RQ1 RNase-Free DNase (Promega) and cDNA was prepared using random hexamer oligonucleotides and Superscript II (Invitrogen). Primers used for qPCR analysis are listed in Table S9.

### Immunofluorescence staining of OSCs

2 days following transfection, cells were plated on concavalin A coated coverslips. After 4 hours, cells were fixed with formaldehyde solution (4% formaldehyde in PBS) for 15 min at room temperature. Fixed cells were washed twice with PBS for 5 min, permeabilized with PBX (0.1 % Triton X-100 in PBS) for 10 min and washed again with PBS for 5 min. Blocking was done in BBS (1 % BSA in PBS) for 30 min and the primary antibody was diluted in BBS and incubated ON at 4°C. Following three washing steps with PBS, the fluorophore-conjugated secondary antibody was diluted in BBS and cells were incubated with it for 1 hour at room temperature in the dark. The stained cells were washed three times with PBS, the second wash containing DAPI. The mounted samples were imaged with a Zeiss LSM-780 confocal microscope and the images were processed using FIJI/ImageJ. Antibodies are listed in Table S11.

### Immunofluorescence staining of ovaries

After dissecting ovaries into ice cold PBS (max 30 min), ovaries were fixed with 4% formaldehyde and 0.3 % Triton X-100 in PBS for 20 min at room temperature. Fixed ovaries were washed 3x with PBX (0.3 % Triton X-100 in PBS) for 10 min and blocked in BBX (1 % BSA and 0.3 % Triton X-100 in PBS) for 30 min. Primary antibody was diluted in BBX and ovaries were incubated with it 24 hours at 4 °C. Following three washing steps with PBX, the fluorophore-conjugated secondary antibody was diluted in BBX and ovaries were incubated with it for ON at 4°C in the dark. The stained ovaries were washed three times with PBX, the second wash containing DAPI. The mounted samples were imaged with a Zeiss LSM-780 confocal microscope and the images were processed using FIJI/ImageJ. Antibodies are listed in Table S11.

### Single molecule RNA Fluorescence In Situ Hybridization (FISH)

*mdg1* RNA FISH on ovaries was performed as described (Mohn et al., 2014) using CAL Fluor Red 590-labeled Stellaris oligo probes (Table S12). After the RNA FISH protocol, egg chambers were blocked with SBX (1% BSA; 0.1% Triton X-100; 2xSSC) for 30 min and then incubated with primary anti-GFP antibody (Abcam) for 24h at 4°C. After 3x washing (10 min with SBX), samples were incubated with fluorescent secondary antibody for 12h at 4°C. Stacks of soma nuclei were imaged on a Zeiss LSM780 confocal microscope and a maximum intensity projection of 3 slices was generated.

### Small RNA-seq

Total RNA was isolated with TRIzol reagent according to the manufacturer’s instructions, and 2S rRNA was depleted as described (Hayashi et al., 2016). Small RNA libraries were generated as described (Jayaprakash et al., 2011). In brief, using radio-labelled oligonucleotides as size-markers, 18 to 29nt long RNAs were purified by PAGE. The 3′ linker (containing four random nucleotides) was ligated with T4 RNA ligase 2, truncated K227Q (NEB) overnight at 16°C. Following PAGE purification, the 5′ linker (containing four random nucleotides) was ligated to the small RNAs using T4 RNA ligase (NEB) overnight at 16°C. After PAGE purification, the linker-ligated RNAs were reverse transcribed and PCR amplified. Sequencing was performed with HiSeq2500 (Illumina) in single-read 50 mode.

### Small RNA-seq analysis

Sequencing reads were trimmed by removal of the adaptor sequences and the four random nucleotides flanking the small RNA. These reads were pre-mapped to the *Drosophila melanogaster* rRNA precursor, the mitochondrial genome and unmapped reads were mapped to the *Drosophila melanogaster* genome (dm6), all using Bowtie (Langmead et al., 2009) (release 1.2.2) with 0 mismatch allowed. Genome mapping reads were intersected with Flybase genome annotations (r6.18) using Bedtools (Quinlan and Hall, 2010) (2.27.1). Reads mapping to rRNA, tRNA, snRNA, snoRNA loci and the mitochondrial genome were removed from the analysis. The quantification of small RNAs was carried out as described in (Andersen et al., 2017). with the following modifications: as a minimal count per 1kb tile cutoff, a value which includes 80% of all reads in the control libraries was used (98 for OSC-KD, 29 for GLKD). Tiles with a mappability below 20% were excluded from the analysis. Annotation groups were based on RefSeq assembly release 6. Tiles overlapping with genes and piRNA clusters were annotated as genic and respective cluster, tiles without annotation were grouped as ‘other’. All sequenced libraries with their GEO Accession number are listed in Table S13.

### RNA-seq with rRNA depletion

We modified the protocol published in (Morlan et al., 2012). Total RNA was isolated with TRIzol reagent, which was further purified by RNAeasy columns with on-column DNase I digest (Qiagen), all according to the manufacturer’s instructions. Depletion of rRNA from the purified total RNA was done by using a mix of antisense oligonucleotides matching *Drosophila melanogaster* rRNAs (listed in Table S14) and the Hybridase Thermostable RNase H (Epicentre) which specifically degrades RNA in RNA-DNA hybrids. The oligonucleotides were added to the RNA in RNase H Buffer (20 mM Tris-HCl pH=8, 100 mM NaCl) and annealed with a temperature gradient from 95 °C to 45 °C. The hybrids were digested at 45 °C for 1 hour. Next, DNA was digested with TURBO DNase (Invitrogen) and RNA was purified using RNA Clean & Concentrator-5 (Zymo) according to the manufacturer’s instructions. Libraries were prepared using a NEBNext Ultra Directional RNA Library Prep Kit for Illumina (NEB) according to the protocol and sequenced on a HiSeq2500 (Illumina) in single-read 50 mode.

### RNA-seq with polyA selection

Total RNA was isolated with TRIzol reagent. Poly(A)+ RNA enrichment was performed with Dynabeads Oligo(dT)^25^ (Thermo Fisher), with two consecutive purifications according to the manufacturer’s instructions. Next, cDNA was prepared using NEBNext Ultra II RNA First and Second Strand Synthesis Module. The cDNA was purified with AmpureXP beads and library was prepared with NEBNext Ultra II DNA Library Prep Kit Illumina (NEB) according to the protocol and sequenced on a HiSeq2500 (Illumina) in single-read 50 mode.

### RNA-seq analysis

Sequencing reads were trimmed by removal of the adaptor sequences. Reads were mapped to the *Drosophila melanogaster* rRNA precursor and the mitochondrial genome using Bowtie (Langmead et al., 2009) (release 1.2.2) with 0 mismatches allowed. Remaining reads were mapped to the *Drosophila melanogaster* genome (dm6) using STAR (Dobin et al., 2013) (v.2.5.2b; settings: --outSAMmode NoQS --readFilesCommand cat --alignEndsType Local --twopassMode Basic -- outReadsUnmapped Fastx --outMultimapperOrder Random -- outSAMtype SAM --outFilterMultimapNmax 1000 -- winAnchorMultimapNmax 2000 --outFilterMismatchNmax 0 -- seedSearchStartLmax 30 --alignSoftClipAtReferenceEnds No -- outFilterType BySJout --alignSJoverhangMin 15 -- alignSJDBoverhangMin 1). Genome mapping reads were intersected with Flybase genome annotations (r6.18) using Bedtools (Quinlan and Hall, 2010) (2.27.1). Reads mapping to rRNA, tRNA, mitoRNA were excluded from further analysis.

### Differential gene expression analysis

Genome matching reads were randomized in order and quantified using Salmon (Patro et al., 2017) (v.0.10.2; settings: -- dumpEqWeights --seqBias --gcBias --useVBOpt -- numBootstraps 100 -l SF --incompatPrior 0.0 -- validateMappings). Salmon results were further processed using wasabi (https://github.com/COMBINE-lab/wasabicommitID=478c133). DGE analysis was performed pairwise between libraries using sleuth (Pimentel et al., 2017) (v0.30.0; settings: extra_bootstrap_summary = TRUE transform_fun_tpm = function(x) log2(x + 0.5), read_bootstrap_tpm = TRUE, gene_mode = TRUE) and running the wald-test function. The sleuth model is a measurement error in the response model. It attempts to segregate the variation due to the inference procedure by Salmon from the variation due to the covariates -- the biological and technical factors of the experiment. For the Wald test, the effect-size represents the estimate of the selected coefficient. It is analogous to, but not equivalent to, the fold-change. The transformed values are on the log2 scale, thus the estimated coefficient is also on the log2 scale. This value takes into account the estimated ‘inferential variance’ estimated from the Salmon bootstraps. For TEs and mRNAs, we required a minimum of TPM >5 in any of the analyzed libraries.

### ChIP-seq

Chromatin immunoprecipitation (ChIP) was carried out according to (Lee et al., 2006), with minor modifications. In brief, OSCs were crosslinked with 1% formaldehyde, quenched with glycine, washed with PBS, collected by centrifugation and pellets were flash-frozen in liquid nitrogen. Chromatin was prepared using Lysis Buffer 1, 2 and 3 from (Lee et al., 2006) and sonication was performed with a Covaris E220 Ultrasonicator for 20 min. For immunoprecipitation, anti H3K9me3 and RNA Pol II antibodies (Table S11), were coupled to Protein G and Protein A Dynabeads, respectively. Sheared chromatin was incubated with the bead-coupled antibodies for 4 hours at 4 °C, beads were washed, and elution plus de-crosslinking was performed at 65 °C overnight. Following RNase A and proteinase K treatment, DNA was purified with ChIP DNA Clean & Concentrator Kit (Zymo). ChIP-qPCR was performed to test the efficiency of the ChIP and libraries were prepared with NEBNext Ultra DNA Library Prep Kit Illumina (NEB) according to the protocol and sequenced on a HiSeq2500 (Illumina) in single-read 50 mode.

### ChIP-seq analysis

Sequencing reads were trimmed by removal of the adaptor sequences and filtered for a minimal length of 18 nucleotides. Reads were mapped to the *Drosophila melanogaster* rRNA precursor, the mitochondrial genome and the genome (dm6) using Bowtie (Langmead et al., 2009) (release 1.2.2), all with 0 (genome wide analysis) or 3 (TE-consensus analysis) mismatches allowed. BigWig files were generated using Homer (Heinz et al., 2010) and UCSC BigWig tools (Kent et al., 2010). Heatmaps and meta profiles were generated with Deeptools within Galaxy using BigWig files. The genomic coordinates of euchromatic TE insertions were determined in (Sienski et al., 2012) and the same Piwi-regulated TEs were used as in (Sienski et al., 2015). To calculate log2 fold change values relative to control knockdown, bigwigCompare was used with a pseudo-count of 1. To determine Piwi-dependent H3K9me3 regions, the quantification of ChIP-seq reads was carried out as described (Andersen et al., 2017) with the following modifications: As a minimal count per tile cutoff a value of 150 reads was used and tiles with a mappability below 20% were excluded from the analysis. Piwi-dependent regions were classified by a log2 fold change > 2 when comparing control knockdown with Piwi knockdown.

For TE consensus analysis, genome mapping reads longer than 23 nucleotides were mapped to TE consensus sequences using STAR (Dobin et al., 2013) (v.2.5.2b; settings: --outSAMmode NoQS -- readFilesCommand cat --alignEndsType Local --twopassMode Basic --outReadsUnmapped Fastx --outMultimapperOrder Random --outSAMtype SAM --outFilterMultimapNmax 1000 -- winAnchorMultimapNmax 2000 --outFilterMismatchNmax 3 -- seedSearchStartLmax 30 --outFilterType BySJout -- alignSJoverhangMin 15 --alignSJDBoverhangMin 1). Multiple mappings were only allowed within one transposon and read-counts were divided equally to the mapping positions. For plotting, read-counts were normalized to 10 million sequenced reads, converted to bedgraph tracks using Bedtools (2.27.1) (Quinlan and Hall, 2010) and plotted in RStudio. All sequenced libraries with their GEO Accession numbers are listed in Table S13.

### Protein co-immunoprecipitation from nuclear OSC cell lysates

OSCs were collected after trypsinization by centrifugation, washed with PBS and centrifuged again. The cell pellet was resuspended in LB1 (10 mM Tris-HCl pH=7.5, 2 mM MgCl_2_, 3 mM CaCl_2_, freshly supplemented with Complete Protease Inhibitor Cocktail (Roche)), incubated at 4°C for 10 min followed by a centrifugation step. The pellet was resuspended in LB2 (10 mM Tris-HCl pH=7.5, 2 mM MgCl_2_, 3 mM CaCl_2_, 0,5 % IGEPAL CA-630, 10 % glycerol, freshly supplemented with Complete Protease Inhibitor Cocktail (Roche)), incubated at 4°C for 10 min followed by a centrifugation step. The isolated nuclei were lysed in LB3 (50 mM Tris-HCl pH=8, 150 mM NaCl, 2 mM MgCl_2_, 0,5 % Triton X-100, 0,25 % IGEPAL CA-630, 10 % glycerol, freshly supplemented with Complete Protease Inhibitor Cocktail (Roche)), incubated at 4°C for 20 min followed by a centrifugation step. Nuclear lysate was used for immunoprecipitation with Flag M2 Magnetic Beads (Sigma) for 2h at 4°C. The beads were washed 3x 10 min with LB3 and were either used for mass spectrometry analysis or the proteins were eluted in 1× SDS buffer with 5 min incubation at 95°C for western blotting.

### Protein co-immunoprecipitation from S2 cell lysates

S2 cells were transfected using Cell Line Nucleofector kit V (Amaxa Biosystems) with the program G-030, using 8 million cells per transfection. S2 cells were co-transfected with FLAG-tagged and GFP-tagged protein encoding plasmids. After two days, cells were collected by centrifugation, washed with PBS and collected again. The cell pellet was resuspended in LB (30 mM Tris-HCl pH=7.5, 150 mM NaCl, 2 mM MgCl_2_, 0,5 % Triton X-100, 10 % glycerol, freshly supplemented with Complete Protease Inhibitor Cocktail (Roche)), incubated at 4°C for 20 min followed by a centrifugation step. The total cell lysate was used for immunoprecipitation with GFP-Trap magnetic beads (ChromoTek) for 2h at 4°C. The beads were washed 3x 10 min with LB and the proteins were eluted in 1× SDS buffer with 5 min incubation at 95°C.

### Western blot

Proteins were separated by SDS–polyacrylamide gel electrophoresis (PAGE) and transferred to a 0.2 μm nitrocellulose membrane (Bio-Rad). The membrane was blocked with 5% milk in PBX (0.05 % Triton X-100 in PBS) and were incubated with primary antibody ON at 4°C. After three washes with PBX, the membrane was incubated with HRP-conjugated secondary antibody for 1h, followed by three PBX washes. The membrane was incubated with Clarity Western ECL Blotting Substrate (Bio-Rad) and imaged with a ChemiDoc MP imaging system (Bio-Rad). Antibodies are listed in Table S11.

### Mass spectrometry analysis

Co-immunoprecipitated proteins coupled to magnetic beads were digested with LysC on the beads, eluted with glycine followed by trypsin digestion. Peptides were analyzed using an UltiMate 3000 RSLCnano System (Thermo Fisher Scientific) coupled to a Q Exactive HF mass spectrometer (Thermo Fisher Scientific), equipped with a Proxeon nanospray source (Thermo Fisher Scientific). Peptides were loaded onto a trap column (Thermo Fisher Scientific, PepMap C18, 5 mm × 300 μm ID, 5 μm particles, 100 Å pore size) at a flow rate of 25 μL/min using 0.1% TFA as mobile phase. After 10 min, the trap column was switched in line with the analytical column (Thermo Fisher Scientific, PepMap C18, 500 mm × 75 μm ID, 2 μm, 100 Å). Peptides were eluted using a flow rate of 230 nl/min and a binary 3h gradient. The gradient starts with the mobile phases: 98% A (water/formic acid, 99.9/0.1, v/v) and 2% B (water/acetonitrile/formic acid, 19.92/80/0.08, v/v/v), increases to 35%B over the next 180 min, followed by a gradient in 5 min to 90%B, stays there for 5 min and decreases in 2 min back to the gradient 98%A and 2%B for equilibration at 30°C.

The Q Exactive HF mass spectrometer was operated in data-dependent mode, using a full scan (m/z range 380-1500, nominal resolution of 60,000, target value 1E6) followed by MS/MS scans of the 10 most abundant ions. MS/MS spectra were acquired using normalized collision energy of 27, isolation width of 1.4 m/z, resolution of 30.000 and the target value was set to 1E5. Precursor ions selected for fragmentation (exclude charge state 1, 7, 8, >8) were put on a dynamic exclusion list for 60 s. Additionally, the minimum AGC target was set to 5E3 and intensity threshold was calculated to be 4.8E4. The peptide match feature was set to preferred and the exclude isotopes feature was enabled.

For peptide identification, the RAW-files were loaded into Proteome Discoverer (version 2.1.0.81, Thermo Scientific). All hereby created MS/MS spectra were searched using MSAmanda v2.1.5.9849, Engine version v2.0.0.9849 (Dorfer et al., 2014). For the first step search the RAW-files were searched against *Drosophila melanogaster* reference translations retrieved from Flybase (dmel_all-translation-r6.13; 21,983 sequences; 20,112,742 residues), using the following search parameters: The peptide mass tolerance was set to ±5 ppm and the fragment mass tolerance to 15ppm. The maximal number of missed cleavages was set to 2. The result was filtered to 1 % FDR on protein level using Percolator algorithm integrated in Thermo Proteome Discoverer. A sub-database was generated for further processing. Peptide areas were quantified using an in-house developed tool APQuant: http://ms.imp.ac.at/index.php?action=peakjuggler (Doblmann et al., 2018).

### Protein expression and purification

The UBA domain of *Drosophila melanogaster* Nxf2 (residues 781-841) and Panoramix helix (residues 311-340) were covalently linked through a KLGSHM linker in one expression cassette. In addition, the NTF2-like domain of *Drosophila melanogaster* Nxf2 (residues 573-777, NTF2l) and full-length Nxt1 (residues 1-133) were cloned into a modified RSFduet-1 vector (Novagen) with an N-terminal His6-SUMO tag on the NTF2-like domain and no tag on Nxt1. Proteins were expressed in *E. coli* strain BL21(DE3) RIL (Stratagene). The cells were grown at 37°C until OD600 reached 0.8, then the media was cooled to 16°C and IPTG (isopropyl β-D-1-thiogalactopyranoside) was added to a final concentration of 0.35 mM to induce protein expression overnight at 16°C. The cells were harvested by centrifugation at 4°C and disrupted by sonication in Binding buffer (20 mM Tris-HCl pH 8.0, 500 mM NaCl, 20 mM imidazole) supplemented with 1 mM PMSF (phenylmethylsulfonyl fluoride) and 3 mM β-mercaptoethanol. After centrifugation, the supernatants were loaded onto 5 ml HisTrap Fastflow column (GE Healthcare). After extensive washing with Binding buffer, the complex was eluted with Binding buffer supplemented with 500 mM imidazole. The His6-SUMO tag was removed by Ulp1 protease digestion during dialysis against Binding buffer and separated by reloading onto HisTrap column. The flow-through fraction was further purified by HiTrap Q FF column and Superdex 75 16/60 column (GE Healthcare). The pooled fractions were concentrated to 35 mg/ml (UBA-linker-helix) and 10 mg/ml in crystallization buffer (20 mM Tris-HCl pH 7.5, 300 mM NaCl, 1 mM DTT).

### Crystallization, data collection and structure determination

As the complex of the UBA domain with the Panoramix helix was not very stable and the yield was very low by co-expression, we covalently linked the UBA domain and the Panoramix helix in one cassette by linkers of different length (KL(GS)_n_HM, n=1, 2, 4, 6). Only the construct with the six-residue KLGSHM linker produced crystals. Crystals of the UBA-linker-helix were grown in 0.095 M sodium citrate pH 5.6, 19% (v/v) Isopropanol, 19% (w/v) PEG 4000, 5% (v/v) glycerol. Crystals of the NTF2-like domain-Nxt1 complex were grown from a solution containing 0.1 M MES pH 6.5, 1.6 M magnesium sulfate using the hanging drop vapor diffusion method at 20°C. For data collection, the crystals were flash frozen (100 K) and collected on NE-CAT beam lines 24ID-C and 24ID-E at the Advanced Photo Source (APS), Argonne National Laboratory. The diffraction data of both, the UBA-linker-Panoramix helix and the NTF2l-Nxt1 complex were processed with the iMosfilm (Battye et al., 2011) and the structures were solved by molecular replacement (MR) in PHENIX (Adams et al., 2002) using the structure of the UBA domain of human NXF1/TAP in complex with FxFG peptide (PDB ID: 1OAI) (Grant et al., 2003) and the structure of the human NXF1/TAP(NTF2l)-NXT1 complex (PDB ID: 1JKG) (Fribourg et al., 2001) as search templates. The automatic model building was carried out using the program PHENIX AutoBuild (Adams et al., 2002). The resulting model was refined by PHENIX refinement (Adams et al., 2002) and Refmac5 (Murshudov et al., 1997), and completed manually using COOT (Emsley et al., 2010). The statistics of the diffraction and refinement data are summarized in Table S6. All the molecular graphics were generated with the PyMOL program (https://pymol.org/2/)

### GST pull-down assay

The C-terminal fragment of *Drosophila melanogaster* Nup358 (residues 2395-2426), which contains three FG-repeats, was fused to GST. 40 μg of GST-FG-repeat were incubated with 50 μL Glutathione Sepharose 4B beads (GE healthcare) at 4°C for 3 h. The beads were washed twice with pull-down buffer (20 mM Tris pH 7.5, 100 mM NaCl, 2 mM DTT and 2 mg/ml BSA) and then incubated with 100 μg protein complexes for 4 h at 4°C. An additional two washes were applied using pull-down buffer without BSA. Each sample was analyzed with SDS-PAGE followed by Coomassie staining.

### Recombinant protein expression in insect cells

To generate a Panoramix (263-446) / Nxf2 (full length) / Nxt1 (full length) co-expression plasmid, the individual open reading frames were cloned into a modified version of the pACEBac1 vector (Geneva Biotech), in which the expression cassette is flanked by BsaI restriction enzyme sites. Panoramix was cloned with an N-terminal Twin-Strep-tag, Nxf2 with an N-terminal His6-tag and Nxt1 with an N-terminal FLAG-tag. All constructs contained the intact polyhedrin leader sequence which harbors a mutated ATG (ATT) upstream of the actual start codon. A low-level of non-canonical initiation from an upstream ATT site results in low level of slightly larger versions of the proteins. The three expression cassettes were then combined into a single destination vector via Golden Gate cloning (Engler et al., 2008). The resulting plasmid was transposed into the EmBacY bacmid backbone (Trowitzsch et al., 2010) and transfected into *Spodoptera frugiperda* Sf9 cells to generate a single baculovirus expressing all three genes. The resulting virus was used to infect *Trichoplusia ni* High5 cells at a density of 1 × 10^6^ cells/ml and expression was performed at 21°C. The cells were harvested 4 days after growth arrest, approximately 96-120 hours after infection, collected by centrifugation and the cell pellets were flash-frozen in liquid nitrogen.

### Affinity purification with Twin-Strep-tag

High5 cells were lysed in lysis buffer (LB) (50 mM Tris-HCl pH=8, 150 mM NaCl, 0,05 % TX100, 1mM DDT) freshly supplemented with Complete Protease Inhibitor Cocktail (Roche) and with Benzonase (~10U/ml) for 30 min at 4°C and the lysate was cleared by centrifugation. For purification, a StrepTactin Superflow HC resin (IBA GmbH) was used with the AKTA Purifier FPLC system and the column was equilibrated with 2 column volumes of LB before sample loading. The bound protein complex was eluted with LB supplemented with 5 mM desthiobiotin and analyzed by SDS-PAGE and InstantBlue (Expedeon) staining.

### Size exclusion chromatography (SEC)

After affinity purification, the complex containing fractions were pooled and further purified by SEC using a Superdex 200 10/300 (GE Healthcare) with the AKTA Purifier FPLC system in SEC buffer (SB) (50 mM Tris-HCl pH=8, 150 mM NaCl, 1mM DDT). The column was equilibrated with 2 column volumes of SB before sample loading. The purified complex was analyzed by SDS-PAGE and Coomassie staining.

### Size exclusion chromatography with in-line multi-angle light scattering (SEC-MALS)

SEC-MALS experiments were performed by using an Äkta-MALS system. Proteins (500 μl) at a concentration of 1.5 mg/ml were loaded on Superdex 75 10/300 GL column (GE Healthcare) and eluted with HEPES buffer (20 mM HEPES, pH=7.5, 200 mM NaCl) at a flow rate of 0.2 ml/min. The light scattering was monitored by a miniDAWN TREOS system (Wyatt Technologies) and concentration was measured by an Optilab T-rEX differential refractometer (Wyatt Technologies). Molecular masses of proteins were analyzed using the Astra program (Wyatt Technologies).

### Crosslinking-mass spectrometry (XL-MS)

#### Protein crosslinking and digestion

The purified complex was crosslinked with 4-(4,6-dimethoxy-1,3,5-triazin-2-yl)-4-methyl-morpholinium chloride (DMTMM; final concentration 4mM). To identify the optimal crosslinker concentration, the complex was titrated with different concentrations and crosslinking yield was checked with SDS-PAGE. The complex was reacted with the crosslinker for 30 minutes at room temperature. To stop the reaction, Tris-HCl pH=8 was added to a final concentration of 100 mM. Sodium deoxycholate (SDC) was added to a final concentration of 1.5%. Samples were reduced with dithiothreitol (DTT, 10 mM, 30 min, 60°C), alkylated with iodoacetamide (IAA, 15 mM, 30 min at room temperature in the dark) and diluted to 1% SDC. Proteins were digested for 3h using trypsin (protein/enzyme 30:1, 37°C). With the addition of 1% trifluoro acetic acid (TFA), SDC was precipitated and the digest was stopped. The supernatant was decanted and stored until measurement.

#### Size-exclusion-chromatography (SEC) enrichment

The digested samples were enriched for crosslinks (XLs) prior to LC-MS/MS analysis using SEC. Therefore, approx. 15 µg of the digest was separated on a TSKgel SuperSW2000 column (300 mm × 4.5 mm × 4 μm, Tosoh Bioscience). The three high mass fractions were collected and measured on the mass spectrometer.

#### LC-MS/MS

Digested peptides were separated using a Dionex UltiMate 3000 HPLC RSLC nanosystem prior to MS analysis. The HPLC was interfaced with the mass spectrometer via a Nanospray Flex™ ion source. For sample concentrating, washing and desalting, the peptides were trapped on an Acclaim PepMap C-18 precolumn (0.3×5mm, Thermo Fisher Scientific), using a flowrate of 25 µl/min and 100% buffer A (99.9% H2O, 0.1% TFA). The separation was performed on an Acclaim PepMap C-18 column (50 cm x 75 µm, 2 µm particles, 100 Ä pore size, Thermo Fisher Scientific) applying a flowrate of 230 nl/min. For separation, a solvent gradient ranging from 2-35% buffer B (80% ACN, 19.92% H2O, 0.08% TFA) was applied. The applied gradient varied from 60-90 min, depending on the sample complexity.

#### Mass Spectrum Acquisition

MS1 spectra were recorded at a resolution of 120000 ranging from 350-1600 m/z (AGC 1e6, 60 ms max. injection time). The top 10 most intense ions from MS1 were selected for fragmentation. MS2 spectra were recorded at 30,000 resolution (AGC 5e4, max. injection time 150 ms, isolation width 1.0 m/z). DMTMM crosslinks were fragmented with higher energy C-trap dissociation (HCD) using a stepped collision energy of 30-33-35%. Once a precursor was selected for MS2, it was excluded from fragmentation for 30sec.

#### Data Analysis

Raw files were analyzed with pLink (Fan et al., 2015) (Version 2.3.3) using the settings as described above. Used crosslinker: DMTMM (−18.0116 Da, reactivity towards lysine, protein N-terminus, serine, threonine and tyrosine or aspartate, glutamate and the protein C-terminus, respectively); MS1 accuracy: 10 ppm; MS2 accuracy: 20 ppm; used enzyme: trypsin; max. missed cleavages: 4; minimum peptide length: 5; max. modifications: 4; static modifications: carbamidomethylation (cysteine, +57.021 Da); dynamic modifications: oxidation (methionine, +15.995 Da). For the database search a database containing the three crosslinked proteins was used and the false discovery rate (FDR) was set to 1%. To reduce the number of false positives, XLs were manually validated. For XL visualization, xiNET was used (Combe et al., 2015).

### Recombinant protein expression in bacterial cells

The open reading frame encoding the Nxf2(1-284) fragment or its point mutant variant was cloned into pET21a with an N-terminal GB1 solubility enhancing tag. Protein expression was in BL21DE3 cells, which were grown at 37°C until OD600=0.6-0.8 and then induced with 0.1 mM IPTG for 18 hours at 18°C. The cell pellet was resuspended in lysis buffer (50mM NaPO_4_ pH=8, 150 mM NaCl, 0.1 % Triton X-100, 10 mM imidazole, 10% glycerol, 5 mM 2-mercaptoethanol, freshly supplemented with 1 mM PMSF and Complete Protease Inhibitor Cocktail (Roche). Lysozyme (10 mg/ml) was added and the cell suspension was incubated for 30 min at 4°C and then sonicated for 15 min at 40% power output with 30% duty cycle (3 sec ON / 7 sec OFF). The sonicated suspension was centrifuged for 20 min at 19,000 g at 4°C. The supernatant was loaded onto pre-equilibrated TALON Metal Affinity Resin (Takara) and incubated for 2 hours. The resin was washed with 20 x bed volume wash buffer (50 mM Tris-HCl pH=7.25, 500 mM NaCl, 20 mM imidazole, 0.1 % Triton X-100, 10 % glycerol, 5 mM 2-mercaptoethanol). The proteins were eluted with 5 x bed volume elution buffer (50 mM Tris-HCl pH=8 300 mM NaCl, 300 mM imidazole, 0.1 % Triton X-100, 10 % Glycerol) and dialyzed against PBS supplemented with 10 % glycerol and 5 mM 2-mercaptoethanol. Post dialysis the protein was aliquoted, flash-frozen in liquid nitrogen and stored at – 80°C for further analysis.

### Electrophoretic mobility shift assay (EMSA)

10 nM [32P] 5’-labelled single-stranded 35 nt RNA (CUCAUCUUGGUCGUACGCGGAAUAGUUUAAACUGU) was incubated with various concentrations of recombinant protein in 10 µl total volume with EMSA binding buffer (10 mM Tris-HCl pH=7.9, 2 mM MgCl_2_, 0.1 mM EDTA, 4% glycerol, 50 mM KCl, 1 mM DTT, 10 µg/ml BSA) for 20 min at 4°C. 2 µl of EMSA loading buffer (50% glycerol, 0.075% bromophenol blue) was added to the samples which were analyzed by 4.8 % PAGE gel in 0.5 x TBE. The radioactive bands were visualized with a Phosphorimager.

### Alpha helix characterization

To determine the physicochemical properties of the predicted alpha-helix within Panoramix, the HeliQuest web server (Gautier et al., 2008) was used. Using an 18 amino acid sliding window (corresponding to a complete helical wheel), the tool predicts hydrophobic surfaces and shows the sequence in a helical wheel representation.

### Phylogenetic tree, orthologue identification and multiple sequence alignment

For phylogenetic reconstruction, Nxf family members from a set of species was extracted, aligned using mafft (v7.407) and the obtained protein sequence alignment was converted to a codon alignment using pal2nal (v14). From this alignment a maximum-likelihood tree was inferred with iqtree (v1.6.7) using best-fitting codon model selection by Modelfinder and 1000 ultrafast bootstrap replicates; visualization using iTol (v4.2.3).

For Nxf protein family collection, proteins showing significant sequence similarity to the NTF2-like domain of *Drosophila melanogaster* Sbr (Nxf1) were collected from the NCBI non-redundant protein database (NCBI nr) using blastp (query NP_524660.1:372-531; species filter: Arthropoda and selected other organisms). Hits were retained if they also showed reciprocal best blast hits to one of the 4 known *Drosophila melanogaster* Nxf family members in blastp searches against the *Drosophila melanogaster* proteome (PTHR10662 members: Nxf3/ FBgn0263232, nxf2/ FBgn0036640, nxf4/ FBgn0051501, sbr/ FBgn0003321). The obtained Nxf protein family set was supplemented with *Drosophila melanogaster* nxf4 protein and its orthologs identified in reciprocal-best-blast-procedure.

For UBA protein domain alignments, a subset of representative Nxf2 protein sequences was selected using an 80% identity cutoff over the C-terminal region (corresponding to NP_524111.3: 726-841). The species selection was also used in alignments of the Nxf1/Tap ortholog groups.

Panoramix ortholog identification was performed using reciprocal psi-blast searches against NCBI nr, using the region of highest conservation among Drosophilid Panoramix proteins as a query (NP_611576.1:292-510). Multiple sequence alignments were visualized in Jalview (v2.10.4), and the secondary structure was predicted with JPRED. Sequence accessions from the alignments are listed in Table S7.

### Data availability statement

All sequencing data used for this study (Table S13) have been deposited with NCBI GEO (accession code GSE120617). The mass spectrometry data have been deposited to the ProteomeXchange Consortium via the PRIDE (Vizcaino et al., 2016) (partner repository under data set identifier PXD011201). All fly lines used in this study are available from the VDRC (http://stockcenter.vdrc.at/control/main). Source data for all gel images are provided as Supplementary Information. Coordinate and structure factors of the UBA-linker-helix and the dmNxf2-Nxt1 complex have been deposited in the Protein Data Bank (PDB) under the accession number 6OPF and 6MRK, respectively.

### Code availability statement

All custom code is based on the publicly available code used in (Andersen et al., 2017) with modifications indicated in the Methods section.

