## Supplementary Figures for "The nascent RNA binding complex SFiNX licenses piRNA-guided heterochromatin formation"

### Supplementary Figure 1

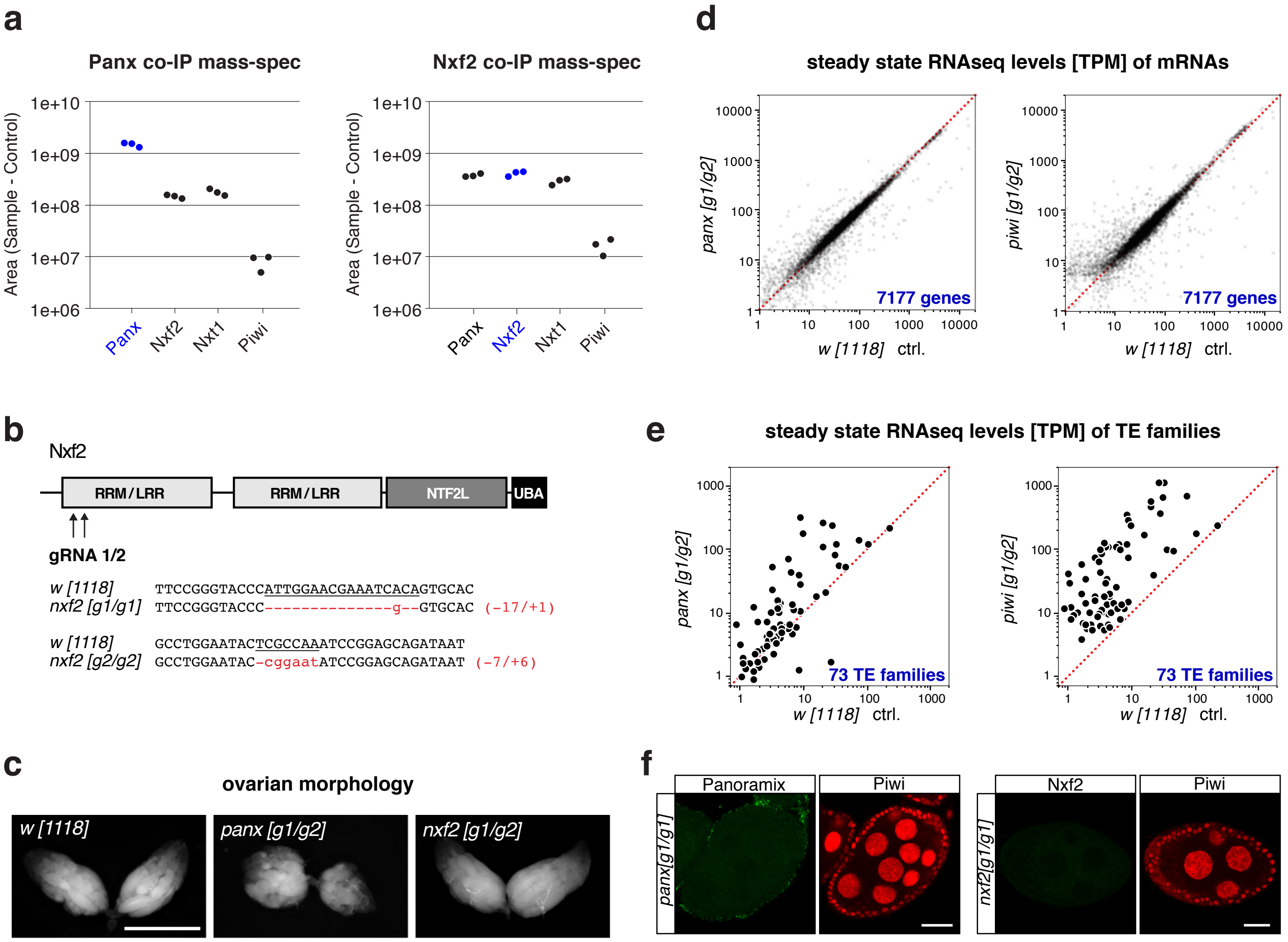


**Supplementary Figure 1. Related to Figure 1**

a, Average peptide peak intensities for indicated proteins in Panoramix or Nxf2 immuno-precipitates (data from experiment shown in Fig. 1a; respective peptide intensities measured in control samples were subtracted).

b, Cartoon depicting frameshift positions caused by the guide RNA-induced insertions/deletions in the Nxf2 protein.

c, Morphology of representative ovaries from flies of indicated genotypes (scale bar: 1mm).

d, e, Scatter plots showing steady state ovarian RNA levels (transcripts per million) of genes (d) and transposons (e) in panx or piwi mutants compared to control.

f, Confocal images of egg chambers (scale bar: 20 µm) from panoramix (left panel) or nxf2 (right panel) mutant flies co-stained for Panoramix or Nxf2, respectively, with Piwi. The remaining staining in the panoramix mutant egg chambers corresponds to a background staining in the surrounding muscle sheet.

### Supplementary Figure 2

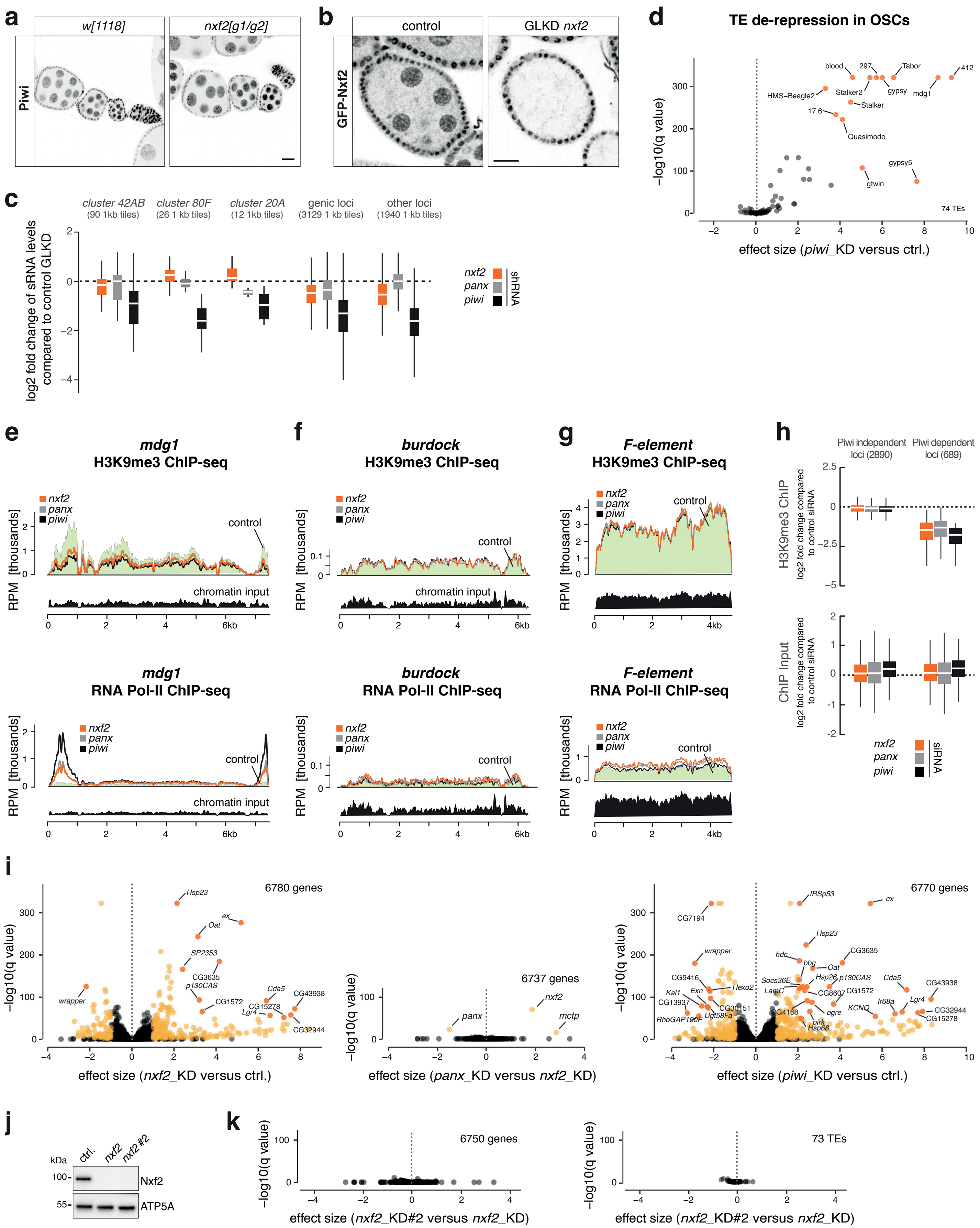

**Supplementary Figure 2. Related to Figure 2**

a, Confocal images of ovarioles (scale bar: 20 µm) from w[1118] control and nxf2 mutant flies stained for Piwi (inverted grayscale).

b, Confocal images showing egg chambers (scale bar: 20 µm) from flies expressing GFP-tagged Nxf2 in addition to indicated germline-specific gene knockdowns (inverted grayscale).

c, Box plot showing fold change (compared to control knockdown) of piRNA levels in ovaries depleted of indicated genes specifically in the germline (box plot definition as in Fig. 2b).

d, Volcano plot showing changes (as effect size) in steady state transposon levels from RNA-seq experiments (n=3). The plot compares piwi knockdown versus control knockdown.

e-g, Density profiles showing normalized reads from H3K9me3 ChIP-seq (top), or RNA Pol II ChIP-seq (bottom) experiments mapping to the mdg1 (e), burdock (f), or F-element (g) transposon consensus sequences (knockdowns indicated).

h, Box plots showing fold changes (compared to control knockdown) of H3K9me3 ChIP-seq signal (top) or input DNA (bottom) in 1kb tiles in OSCs depleted of indicated genes. Piwi dependent and independent regions are shown separately (box plot definition as in Fig. 2b).

i, Volcano plots showing differential gene expression analysis from OSC RNA-seq experiments (gene knockdowns indicated, n=3).

j, Western blots showing knockdown efficiencies of two different nxf2 siRNAs in OSCs. ATP5A served as loading control.

k, Volcano plots showing differential expression analysis of genes (left) and transposons (right) from OSC RNA-seq experiments (gene knockdowns indicated; n=3). The plots compare the two tested independent siRNAs targeting nxf2.

### Supplementary Figure 3

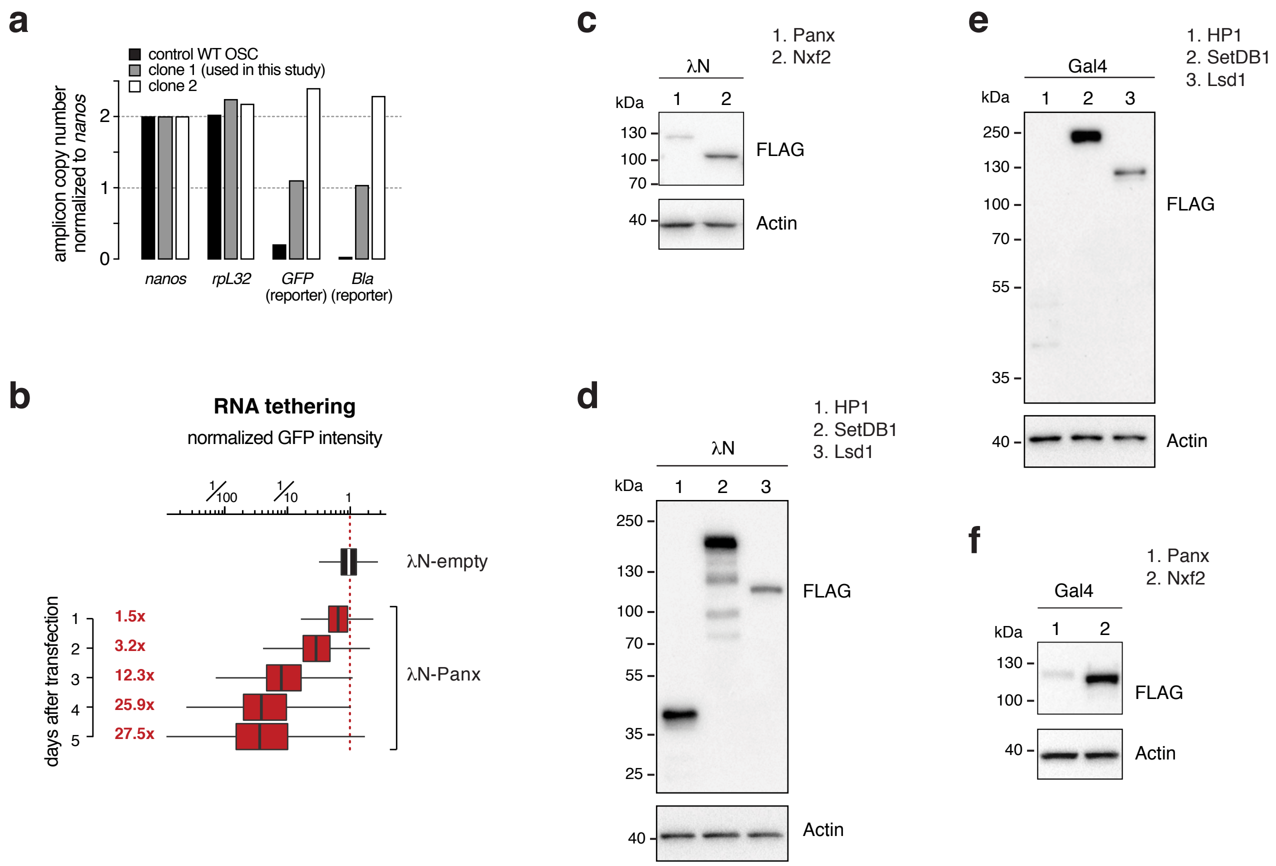


**Supplementary Figure 3. Related to Figure 3**
a, Bar graph showing genomic copy number of indicated genes based on digital droplet PCR. Two clonal reporter OSC lines were analyzed and clone 1, which harbors a single insertion, was used throughout the study.
b, Box plots showing the λN-control normalized GFP intensity in OSCs at indicated days after transfection of λN-tagged Panoramix protein; n=2500 cells; box plot definition as in Fig. 2b).
c-f, Western blots showing levels of indicated FLAG-tagged fusion proteins (λN or Gal4) in OSC lysates (related to Fig. 3c, f, g, h; Actin served as loading control).

### Supplementary Figure 4

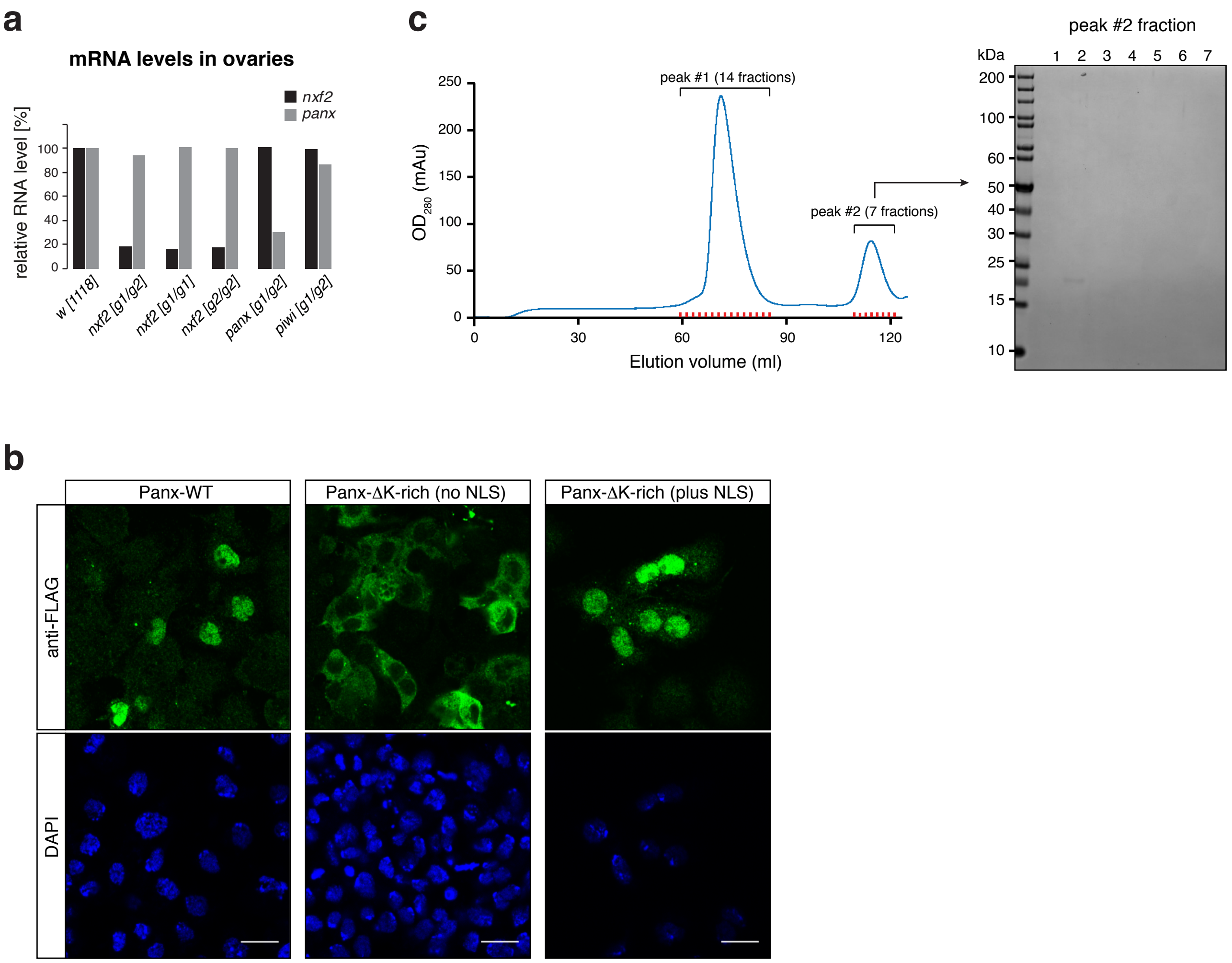


**Supplementary Figure 4. Related to Figure 4**

a, Bar graph showing fold change of RNA-seq levels of indicated genes in mutant ovaries compared to control ovaries (n = 1).

b, Confocal images showing OSCs (scale bar: 10 µm) with indicated, transiently transfected FLAG-tagged Panoramix constructs.

c, Left: Shown is the entire size exclusion chromatogram (Fig. 4f) of the affinity-purified Strep-Panoramix eluate (mAU = milli-absorbance unit). To the right, an SDS-PAGE of the 7 peak fractions from the second detected peak is shown (Coomassie blue staining), indicating that this peak does not contain protein.

**Supplementary Figure 5. Related to Figure 5**

a-c, Western blot analysis of GFP or GFP-Nxf2 immuno-precipitation experiments using lysate from S2 cells transiently co-transfected with indicated FLAG-Panoramix expressing plasmids (relative amount loaded in immunoprecipitation lanes: 3x).

d, Protein sequence alignment of Panoramix (308-446) where the experimentally tested residues are marked in blue. Shown below are predicted secondary structural elements and the conservation score for each position.

e, Helical wheel representation of the predicted amphipathic $\alpha$-helix (322-339) within Panoramix. The point mutations introduced to abolish the Nxf2 interaction are indicated in purple color.

f, Western blot analysis of lysate from S2 cells transiently transfected with indicated FLAG-Panoramix expressing plasmids. Actin served as loading control.

g, Box plots showing GFP intensity in S2 cells 2 days after transfection with plasmids expressing indicated fusion proteins or empty control (numbers indicate fold-change in median GFP intensity normalized to GFP-only control; box plot definition as in Fig. 2b).

h, Western blot analysis of immuno-precipitation experiments (bait: stabilized degron mutant GFP-Panoramix) using lysate from S2 cells transiently co-transfected with indicated FLAG-Nxf2 expressing plasmids (relative amount loaded in immunoprecipitation lanes: 3x).

i, Co-purification of His-SUMO-Panoramix helix with untagged NXF2 UBA domain without linker by two rounds of Ni-NTA affinity purification.

### Supplementary Figure 5

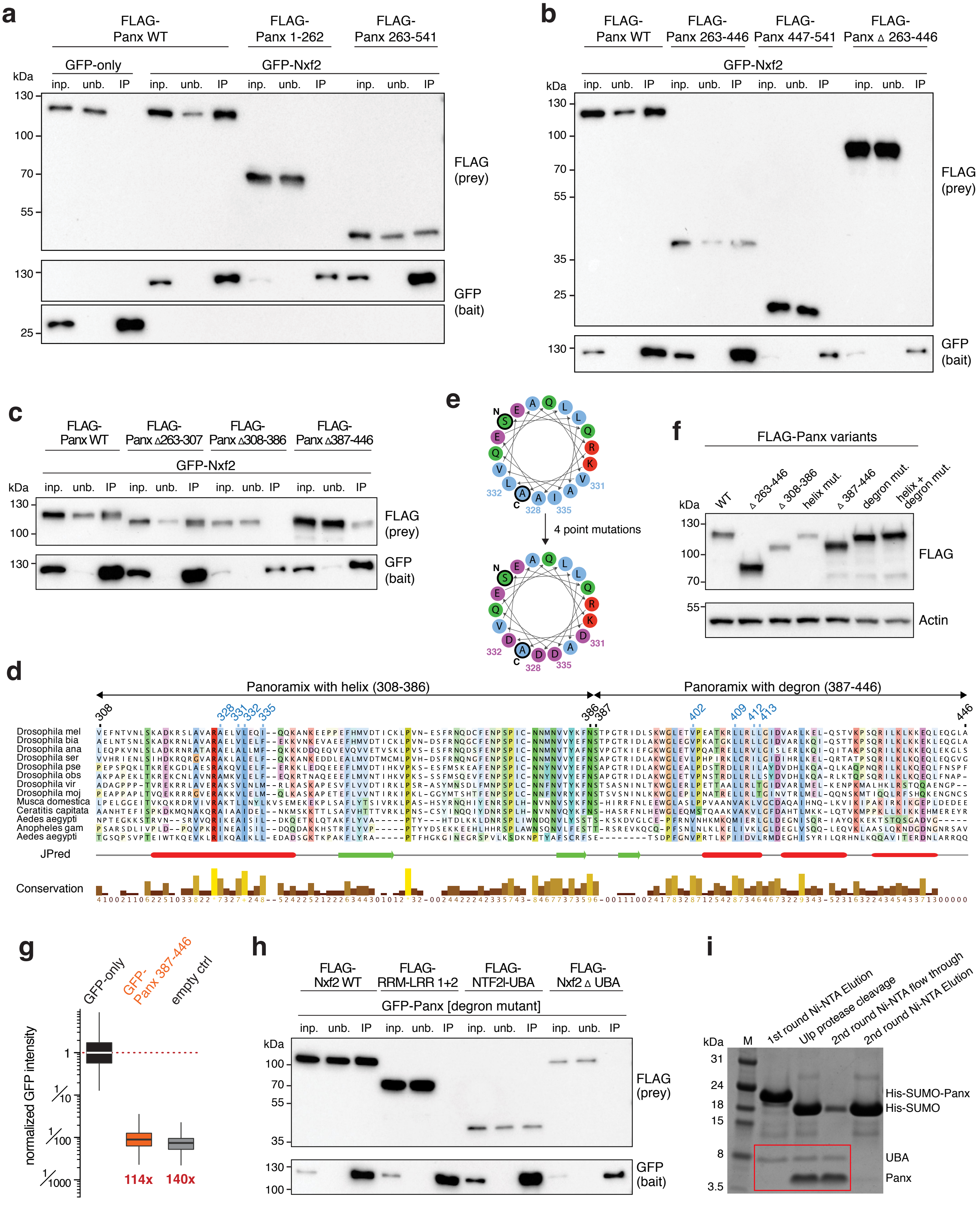

### Supplementary Figure 6

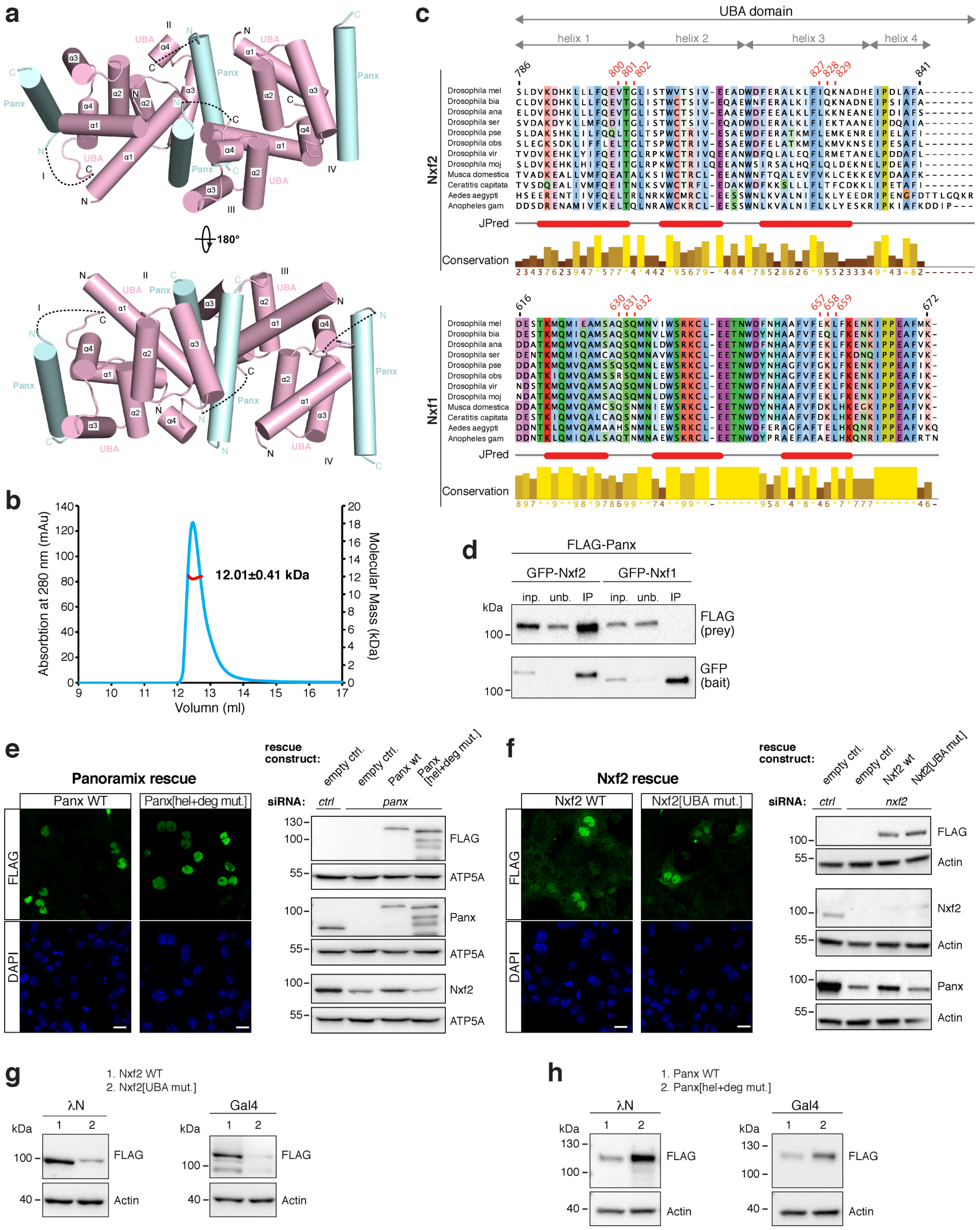

**Supplementary Figure 6. Related to Figure 5**

a, Cartoon view of four UBA-linker-helix monomers in the crystal asymmetric unit. The linkers between the UBA domain and the Panoramix helix are largely invisible and are shown as dashed lines.

b, Molecular weight of UBA-linker-Panoramix helix measured by SEC-MALS assay. The red line represents the SEC-MALS calculated molecular weight, which is 12.0±0.4 kDa. The theoretical molecular weight is 11.1 kDa, indicating that UBA-linker-Panoramix helix is a monomer in solution.

c, Protein sequence alignment of the Nxf2 and Nxf1/Tap UBA domains from indicated insect species. The relevant mutated residues are marked in red. Shown below are predicted secondary structural elements and the conservation score for each position.

d, Western blot analysis of GFP-Nxf2 or GFP-Nxf1 immuno-precipitation experiments using lysate from S2 cells transiently co-transfected with indicated FLAG-Panoramix expressing plasmids (relative amount loaded in immunoprecipitation lanes: 3x).

e, Left: Confocal images showing OSCs (scale bar: 10 µm) with indicated, transiently transfected FLAG-tagged Panoramix constructs. Right: Western blots showing levels of indicated proteins in OSC lysates with indicated knockdowns and transiently transfected FLAG-tagged Panoramix constructs (ATP5A served as loading control).

f, Left: Confocal images showing OSCs (scale bar: 10 µm) with indicated, transiently transfected FLAG-tagged Nxf2 constructs. Right: Western blots showing levels of indicated proteins in OSC lysates with indicated knockdowns and transiently transfected FLAG-tagged Nxf2 constructs (ATP5A served as loading control).

g, h, Western blots showing levels of indicated fusion proteins (left: λN, right: Gal4) in lysates of transiently transfected OSCs (related to Fig. 5i, j; Actin served as loading control).

### Supplementary Figure 7

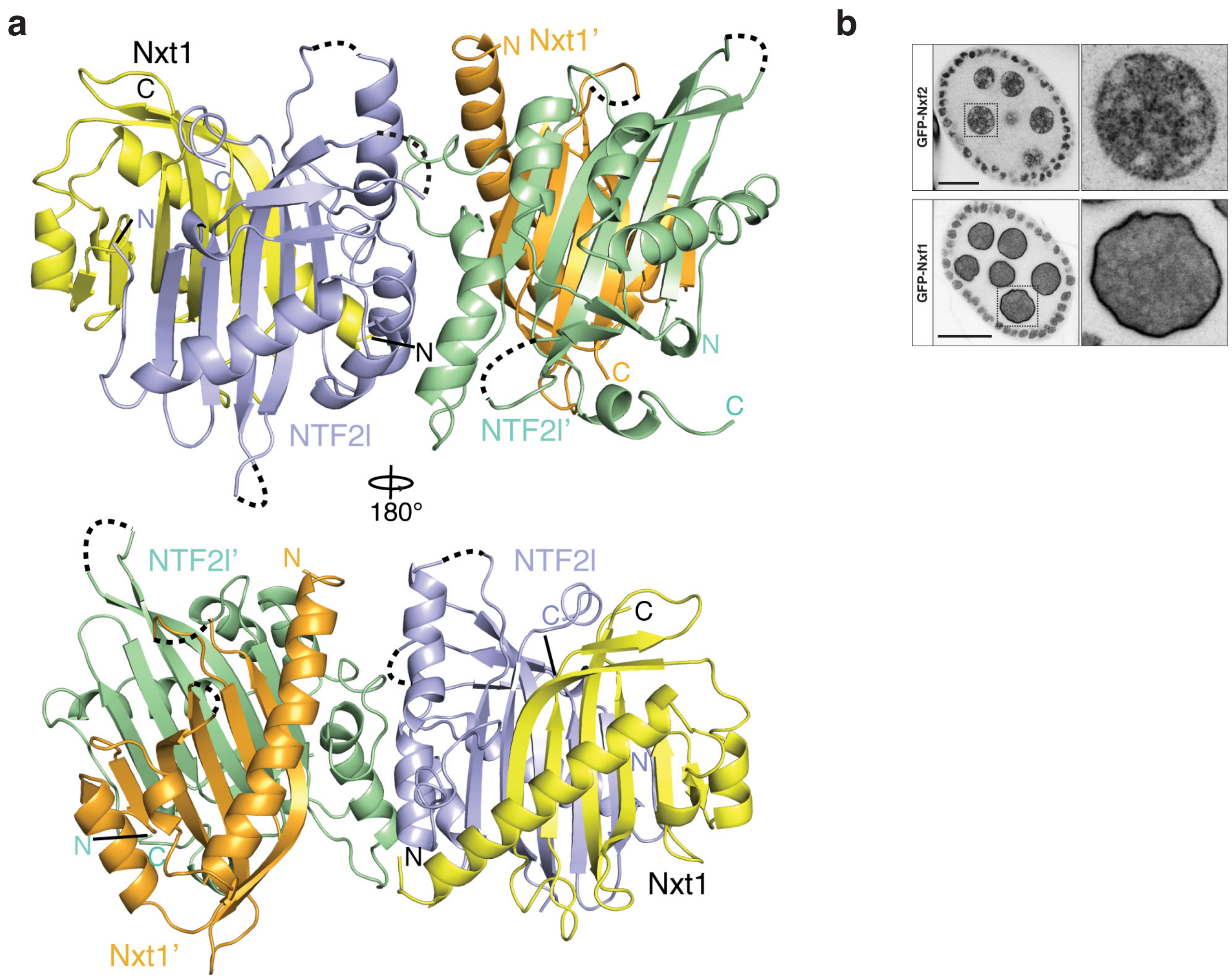


**Supplementary Figure 7. Related to Figure 6**

a, Front and back views of the crystal structure of dmNxf2's NTF2-like domain (purple and green) in complex with dmNxt1/p15 (yellow and orange). Two NTF2-like domain-Nxt1/p15 heterodimers were observed in the crystal asymmetric unit due to crystal packing. Invisible loops in the structure are shown as dashed curves.

b, Confocal images showing egg chambers (left) and individual nurse cell nuclei (right) from flies expressing GFP-tagged Nxf2 or GFP-tagged Nxf1 (scale bar: 20 µm).

### Supplementary Figure 8

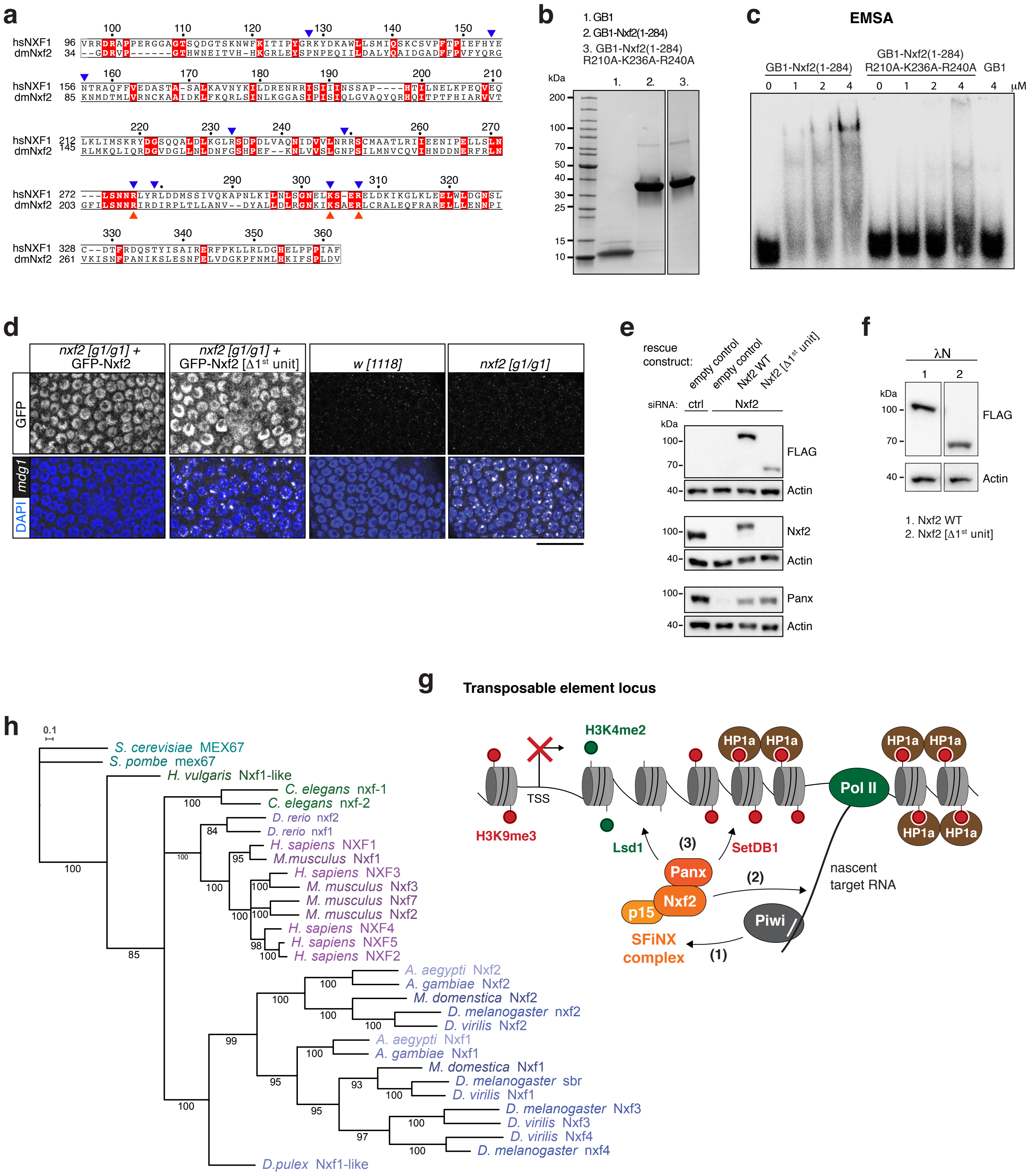

**Supplementary Figure 8. Related to Figure 7**

a, Protein sequence alignment of the RRM/LRR domain of hsNXF1/TAP and the first RRM/LRR domain of dmNxf2. Identical residues are highlighted by red squares. The key hsNXF1/TAP residues involved in CTE RNA binding are indicated by blue triangles; red triangles mark residues which were mutated in dmNxf2 for the EMSA assay in c.

b, Coomassie stained SDS-PAGE showing the recombinant RNA-binding unit #1 of Nxf2, its point mutant variant, and the GB1 control peptide.

c, Phosphorimage showing an Electrophoretic Mobility Shift Assay (EMSA) with labeled single stranded RNA (ssRNA) and increasing amount of the purified Nxf2 protein, its point mutant variant, and the GB1 control peptide.

d, Confocal images depicting somatic cells of egg chambers (scale bar: 20 µm) from flies with indicated genotypes. Expression of the rescue transgenes was tested by GFP fluorescence, transposon derepression was assessed by mdg1 FISH.

e, Western blots showing levels of indicated proteins in OSC lysates with indicated knockdowns and transiently transfected rescue constructs (relates to experiment shown in Fig. 7e; ATP5A served as loading control).

f, Western blots showing levels of indicated λN-fusion proteins in OSC lysates (related to Fig. 7g; Actin served as loading control).

g, Schematic model for the mechanism of Piwi-mediated co-transcriptional silencing utilizing the SFiNX complex. Upon target binding, Piwi interacts with the adaptor complex, SFiNX (1). Within SFiNX, Nxf2 links to the nascent target RNA, possibly after a regulatory switch (2). This allows Panoramix to recruit chromatin effectors, including Lsd1 to demethylate H3K4me2 and SetDB1 to establish H3K9me3, which is then bound by HP1a (3). (TSS: transcription start site)

h, Maximum likelihood phylogenetic tree of NXF sequences from indicated species inferred with iqtree. Numbers represent ultrafast bootstrap branch support values. Scale bar indicates expected number of substitutions per codon site.
